# Vaccines combining slow delivery and follicle targeting of antigens increase germinal center B cell clonal diversity and clonal expansion

**DOI:** 10.1101/2024.08.19.608655

**Authors:** Kristen A. Rodrigues, Yiming J. Zhang, Aereas Aung, Duncan M. Morgan, Laura Maiorino, Parisa Yousefpour, Grace Gibson, Gabriel Ozorowski, Justin R. Gregory, Parastoo Amlashi, Maureen Buckley, Andrew B. Ward, William R. Schief, J. Christopher Love, Darrell J. Irvine

## Abstract

Vaccines incorporating slow delivery, multivalent antigen display, or immunomodulation through adjuvants have an important role to play in shaping the humoral immune response. Here we analyzed mechanisms of action of a clinically relevant combination adjuvant strategy, where phosphoserine (pSer)-tagged immunogens bound to aluminum hydroxide (alum) adjuvant (promoting prolonged antigen delivery to draining lymph nodes) are combined with a potent saponin nanoparticle adjuvant termed SMNP (which alters lymph flow and antigen entry into lymph nodes). When employed with a stabilized HIV Env trimer antigen in mice, this combined adjuvant approach promoted substantial enhancements in germinal center (GC) and antibody responses relative to either adjuvant alone. Using scRNA-seq and scBCR-seq, we found that the alum-pSer/SMNP combination both increased the diversity of GC B cell clones and increased GC B cell clonal expansion, coincident with increases in the expression of *Myc* and the proportion of S-phase GC B cells. To gain insight into the source of these changes in the GC response, we analyzed antigen biodistribution and structural integrity in draining lymph nodes and found that the combination adjuvant approach, but not alum-pSer delivery or SMNP alone, promoted accumulation of highly intact antigen on follicular dendritic cells, reflecting an integration of the slow antigen delivery and altered lymph node uptake effects of these two adjuvants. These results demonstrate how adjuvants with complementary mechanisms of action impacting vaccine biodistribution and kinetics can synergize to enhance humoral immunity.

## INTRODUCTION

Vaccination elicits robust and targeted protection against infection by prompting the immune system to recognize potential threats (*1*). Most licensed vaccines are thought to protect through antibody responses (*1, 2*), whereby antigen-specific helper T cells and B cells are activated and work together in germinal centers (GCs) to generate high-affinity antibody-secreting plasma cells and memory B cells (*3*). Despite the success of vaccination-induced immunity against many pathogens, a number of major challenges remain, such as the development of effective vaccines against HIV and tuberculosis, “universal” vaccines for influenza that could provide cross-seasonal protection, or pan-coronavirus vaccines (*4–7*).

HIV serves as a useful exemplar of challenges common to these “difficult” vaccine cases: A protective vaccine will likely need to elicit several classes of broadly neutralizing antibodies (bnAbs), which recognize conserved sites on the viral envelope across the diversity of circulating viral strains. HIV-infected humans can generate bnAbs, and many classes of bnAbs have been isolated from patients (*8, 9*). However, HIV bnAbs have uncommon features such as extensive somatic hypermutation (SHM), improbable mutations, and very long CDR3 junction lengths (*10, 11*). Consequently, bnAb-precursor B cells are typically rare and present at very low frequencies in the human B cell repertoire (*12–14*).

To overcome these challenges, vaccine regimens capable of recruiting rare B cell clones into the GC reaction and promoting their expansion and affinity maturation may be required. One strategy to modulate the GC response is by manipulating vaccine kinetics, i.e. the timing of inflammatory cue or antigen delivery to draining lymph nodes (dLNs). For example, sustained vaccine delivery over a few weeks using repeated injections or implantable osmotic pumps has been shown to increase the number of unique clones recruited to GCs and greatly increase the size of the GC response compared to traditional bolus vaccine administration (*15–17*). To make this approach more clinically translatable, we converted the most common clinical adjuvant, aluminum hydroxide (alum), into a slow-delivery vehicle by modifying immunogens with short phosphoserine (pSer) peptide tags (*18–20*). Through a ligand exchange reaction between phosphate and hydroxyls, these pSer tags anchor antigens to the surface of alum particles. This approach, which we refer to hereafter as “alum-pSer”, promotes stable retention of the antigen on alum particles *in vivo* and leads to prolonged antigen drainage from the injection site following a bolus injection, which translated into improved GC B cell and serum IgG antibody responses and the development of long-lived bone marrow plasma cells in mice for HIV and SARS-CoV-2 antigens (*18–20*).

A second approach to tune GC responses is via the selection of appropriate adjuvants, which can impact many aspects of the immune response including antigen presentation, immune cell recruitment and retention in dLNs, and inflammatory cytokine production that direct the adaptive immune response (*21, 22*). Saponins are potent adjuvants for promoting humoral response and are used in the licensed Shingrix® and Mosquirix® vaccines from Glaxo-Smith Kline as well as the Novavax SARS-CoV-2 vaccine (*23, 24*). We recently developed a saponin-based adjuvant called SMNP, a ∼40 nm diameter nanoparticle formed by the self-assembly of phospholipids, cholesterol, saponin, and the Toll-like receptor (TLR)-4 agonist monophosphoryl lipid A (MPLA) (*25*). SMNP co-administration has multiple effects on the immune response in both mice and non-human primates, including enhanced lymph trafficking of antigen, increased antigen entry into dLNs, and induction of a cascade of inflammatory cytokines and chemokines in dLNs (*17, 25, 26*). Based on these promising findings, SMNP will shortly enter first-in-human testing through the HIV Vaccine Trials Network (HVTN 144).

Inspired by their complementary mechanisms of action, we previously tested the impact of combining pSer-tagging of antigens for alum anchoring and sustained antigen delivery with co-administration of SMNP. We discovered that this combination adjuvant approach showed striking amplification of humoral responses to both HIV Env and SARS-CoV-2 antigens (*19, 20*): alum-pSer + SMNP immunization led to enhancements in GC B cell and Tfh responses, and increased serum IgG and neutralizing antibody responses. Here, we sought to investigate the immunological basis of these improved humoral immune responses and identify underlying mechanisms that might explain this striking potency of the alum-pSer/SMNP combination. We first performed single-cell RNA sequencing (scRNA-seq) transcriptional profiling and single-cell B cell receptor sequencing (scBCR-seq) of antigen-binding GC B cells from mice immunized with a stabilized HIV Env trimer immunogen termed MD39 combined with alum-pSer, SMNP, or alum-pSer/SMNP adjuvants. These analyses revealed that the combination adjuvant augmented multiple complementary facets of the GC response and increased the proportion of GC B cells in S phase of the cell cycle, suggesting greater antigen acquisition and T cell help. Motivated by these findings, we analyzed the biodistribution of antigen following alum-pSer, SMNP, or combined adjuvant immunization, and discovered that the combination adjuvants uniquely promoted robust accumulation of intact trimer antigen on follicular dendritic cells (FDCs), which persisted for several weeks. These findings indicate that this simple combination adjuvant approach achieves both sustained antigen availability and altered antigen localization that can productively drive important changes in the composition of the GC response, which may be valuable for diverse infectious disease targets.

## RESULTS

### scRNA-seq profiling of antigen-specific GC B cells primed by alum-pSer and SMNP immunizations

To gain insights into how alum-pSer and SMNP impact the humoral immune response, we first carried out a single-cell RNA-seq (scRNA-seq) study of antigen-binding GC B cells elicited by these adjuvants combined with an HIV Env stabilized SOSIP trimer immunogen termed MD39 (*27*). We compared 3 formulations (**Fig. 1A**): For alum-pSer immunization, a peptide tag containing four pSer residues was conjugated to the C-terminus of each MD39 gp140 protomer, leading to three pSer_4_ tags placed at the base of each trimer. When mixed with alum adjuvant, the phosphoserines of these tagged trimers undergo a ligand exchange reaction with hydroxyl groups on the surface of alum, anchoring the trimer immunogen in an oriented fashion to the alum particles (*18–20*) (alum-pSer, **Fig. 1A**). The second vaccine formulation was comprised of MD39 trimer mixed with SMNP adjuvant (SMNP, **Fig. 1A**). SMNP is composed of ∼40 nm particles of saponin, MPLA, lipids, and cholesterol, which self-assemble to form a cage-like structure; SMNP does not interact with the MD39 trimer in solution (**fig. S1**). The third formulation was comprised of the combination of pSer-tagged MD39 bound to alum and mixed with SMNP particles (alum-pSer/SMNP, **Fig. 1A**).

**Fig. 1.**
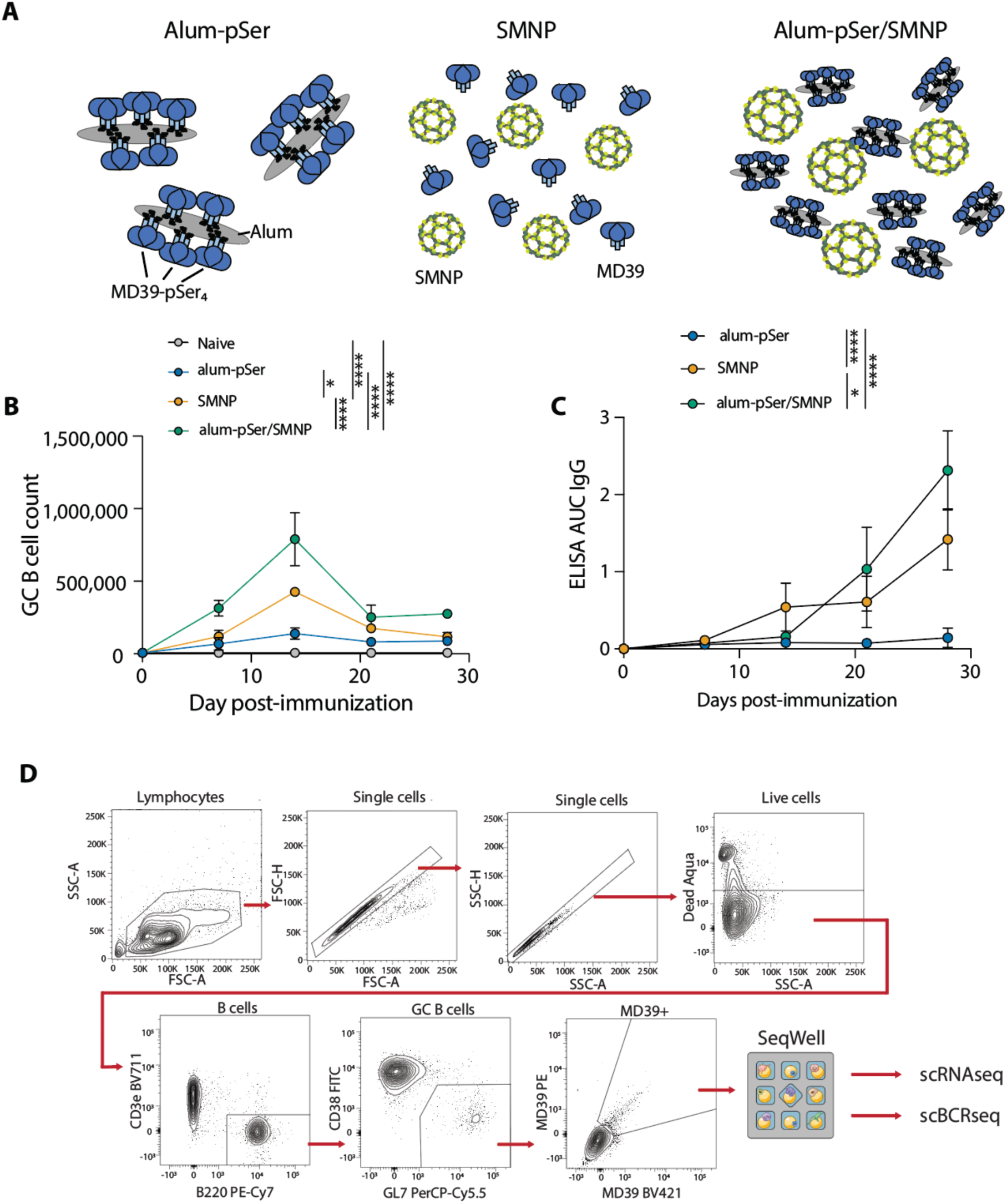
pSer-modified Env trimer anchored on alum combined with SMNP adjuvant amplifies humoral immune responses. (**A**) Schematic of immunization groups. (**B**-**C**) BALB/c mice (*n*=5/group for flow cytometry analysis, *n*=14/group for scRNA-seq and scBCR-seq) were immunized with 5 µg MD39 Env trimer ± 50 µg alum ± 5 µg SMNP. (**B**) Total germinal center (GC) B cell counts over time. Values plotted are mean ± s.e.m. (**C**) Serum IgG antibody responses were assessed longitudinally by ELISA using MD39 captured by lectin. Values plotted are ELISA area under the curve (AUC) mean ± s.d. (**D**) The GC responses in draining lymph nodes were analyzed by flow cytometry, and MD39-binding GC B cells were loaded onto SeqWell arrays for scRNA-seq and scBCR-seq. Statistical significance was determined by two-way ANOVA followed by Tukey’s multiple comparisons test. ns p>0.05, * p<0.05, ** p<0.01, *** p<0.001, **** p<0.0001.

We previously found that MD39 + alum-pSer/SMNP elicited antibody and GC B cell responses substantially superior to either alum-pSer or SMNP alone (*18, 20*). To identify an appropriate timepoint for scRNA-seq analysis, we first immunized BALB/c mice with each formulation and analyzed humoral responses over time (**Fig. 1B-D**). As shown in **Fig. 1B**, GC responses in all 3 groups steadily expanded for two weeks post-immunization, peaking at day 14, and then began contracting. Consistent with our prior findings, there was a clear hierarchy in size of the GC responses, with the combination alum-pSer/SMNP immunization eliciting 1.9-fold and 5.6-fold more GC B cells than SMNP or alum-pSer alone, respectively, at the peak of the response. Serum antibody responses developing over the same time course also showed the strongest response in the combination adjuvant group (**Fig. 1C**). Based on these findings, we carried out scRNA-seq and scBCRseq analyses at the peak of the GC response for each group, day 14 post-immunization. Groups of mice were immunized with each of the 3 vaccine formulations, and the antigen-binding GC B cells were flow sorted for combined scRNA-seq and scBCR-seq using SeqWell, a nanowell-based library preparation (*28–30*) (**Fig. 1D**).

The number of recovered cells in each immunization condition reflected the magnitude of the GC response detected by flow cytometry (**Fig. 2A**). After quality control, we recovered the transcriptome of 11,231 MD39-binding GC B cells, including 149 from alum-pSer, 2608 from SMNP, and 8474 from alum-pSer/SMNP immunized mice (**Fig. 2B**). We first examined the transcriptome data. Leveraging unsupervised clustering and differential gene expression analysis, we identified seven phenotypic clusters (**Fig. 2C-D**). Among them, cluster 1 (C1) showed plasmablast gene signatures such as Cd138 *(Sdc1),* Blimp-1 *(Prdm1), Xbp1, and Ell2* (*31*). C2 upregulated *Ccr6*, *Hhex*, *Fcer2a*, and GC egressing markers *Itga4*, *Itgb7*, *Lmo2*, and *Cmah*, suggesting that C2 cells are likely GC-derived pre-memory B cells (MBCs) (*32–34*). C3 cells expressed genes involved in antigen capturing and presentation (*H2-DMa, H2-Ab1, Ciita, Cr2*) and signaling with T cells (*Cd83, Cd86, Cd40*) (*35, 36*), implicating a light zone (LZ) B cell phenotype. Cell division is the hallmark of dark zone (DZ) B cells (*3, 35*). We demarcated DZ cells into three sub-clusters, where C5 was characterized by S phase genes (*Mcm6, Pcna, Lig1*), C6 by active cycling genes (*Cenpe, Mki67, Cdc20*), and C7 by canonical DZ markers (*Gcsam, Aicda, Cxcr4, and Foxo1*). Lastly, C4 cells showed an intermediate phenotype between LZ and DZ based on their expression of positive-selection and early proliferation markers *Cd40, C1qbp, Mybbp1a, Myc, Mtor, Mif, Bcl2a1b*, and SHM and CSR marker *Ung* (*35, 37–39*), which is also a downstream target of Myc (*40*). Additionally, based on the high expression of Myc– and mTORC1-targeted genes (**fig. S2A**), the majority of C4 cells are likely positively selected B cells. These clusters are consistent with phenotypes observed in prior studies of mouse and human GC B cells (*36, 41–43*) (**fig. S2B-D**).

**Fig. 2.**
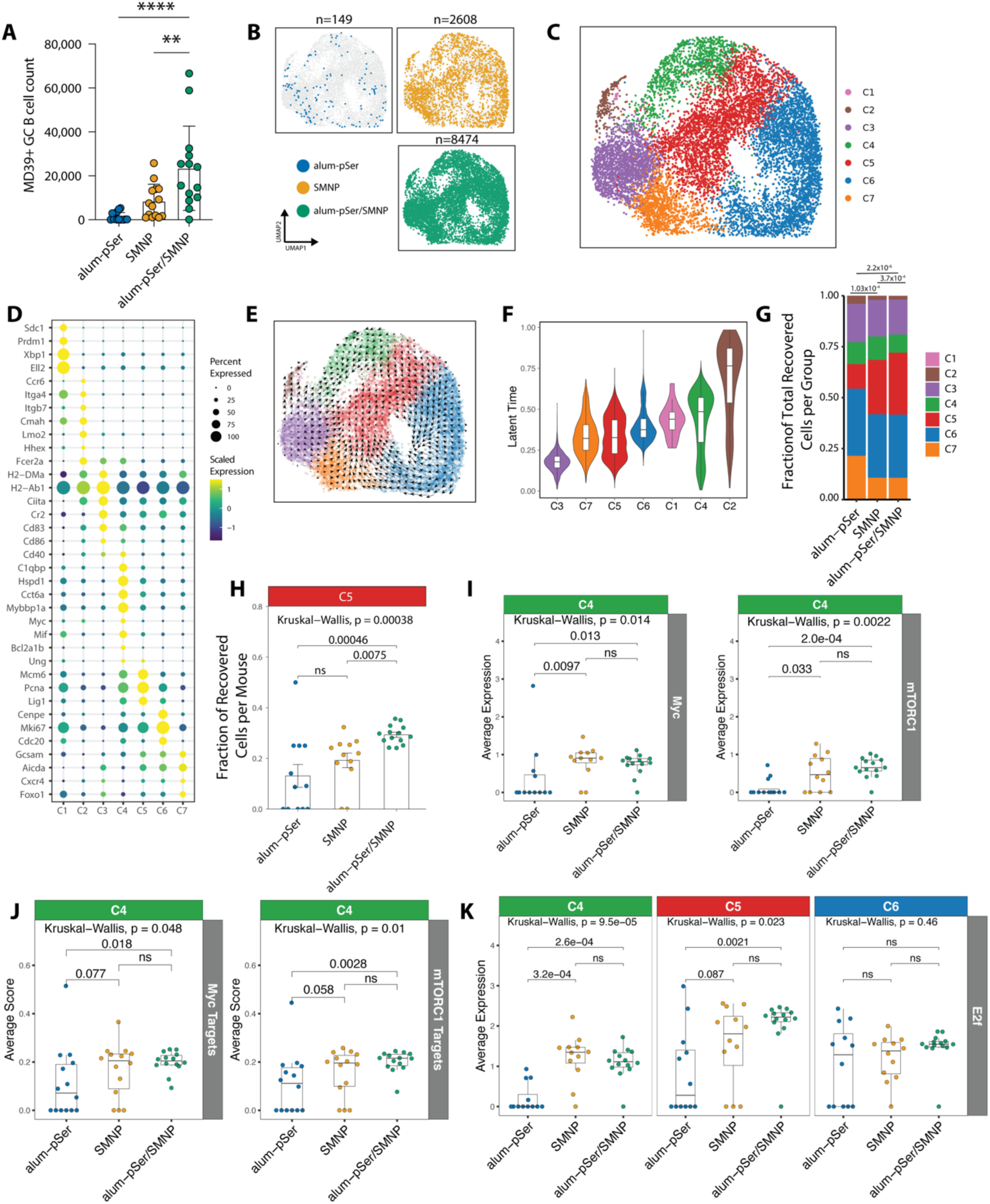
scRNA-seq profiling of MD39-binding germinal center B cells 14 days post-immunization. (**A**) GC B cell counts detected by flow cytometry and loaded onto SeqWell arrays. Shown are mean ± s.e.m. (**B**) UMAP projection of recovered cells by immunization group. Gray dots represent all recovered cells. (**C**) UMAP projection of phenotypic clusters of MD39-binding GC B cells. (**D**) Differentially expressed genes associated with phenotypic clusters. The color indicates scaled expression levels. The dot radius indicates the fraction of cells expressing the gene. (**E**) UMAP projection of RNA velocity vector fields. The length of the arrows indicates the speed of differentiation. (**F**) Latent time distribution by clusters. (**G**) Cluster distribution of recovered cells. (**H**) The fraction of C5 cells per mouse from each vaccine group. (**I**) Average expression of *Myc* and mTORC1 genes (sum of *Mtor, Rptor, Akt1s1,* and *Deptor*) among C4 cells.(**J**) Average module score of *Myc*– and *mTORC1*-target genes among C4 cells. (**K**) Average expression of activating *E2fs*-family transcription factors (sum of *E2f1, E2f2, E2f3*) among C4, C5, and C6 cells. Dots in **H-K** represent the average expression of individual mice. For (**A**), statistical significance was determined by one-way ANOVA followed by Tukey’s multiple comparisons test. ns p>0.05, ** p<0.01, **** p<0.0001. For (**G**), p values are computed with Chi-squared tests with Bonferroni correction. For (**H-K**), p values are computed with Kruskal-Wallis analysis of variance followed by Dunn’s post hoc test.

To further validate our clustering, we performed RNA velocity analysis (**Fig. 2E**). The velocity vector fields showed a bifurcation among LZ cells (C3) towards pre-MBC (C2) or transitioning back to the DZ (C4, C5) (**Fig. 2E**). Cyclical cell division in the DZ was well-reflected by vector fields moving from C5 to C6 to C7 (**Fig. 2E**). The pseudotemporal ordering of antigen-binding GC B cells based on the latent time calculated from RNA velocity revealed a continuum of differentiation trajectory from LZ (C3) or G1 DZ (C7) to cell division in the DZ (C5, C6) and eventual exit as plasmablasts (C1) or pre-MBCs (C2) (**Fig. 2F**, **fig. S2E**). The LZ/DZ intermediate cells (C4) spanned a wide range of latent time, with the majority having high latent time, implicating longer residence in the GC and supporting the posit of positively selected B cells. In summary, our transcriptional profiling is consistent with the current understanding of GC reactions (*3, 35*).

### Alum-pSer/SMNP combination adjuvant elicits an enrichment of S-phase GC B cells

GC B cells recovered from SMNP– and alum-pSer/SMNP-immunized mice showed substantially greater proportions of B cells in S phase (C5, **Fig. 2G-H, fig. S2F**). Such an observation is of interest because the enrichment of S-phase B cells has been correlated with the strength of positive selection occurring in the LZ (*44, 45*). Positively selected LZ B cells express *Myc* and mTOR complex 1 (mTORC1, consisting of *Mtor, Rptor, Akt1s1,* and *Deptor*) as they enter S phase and migrate back to the DZ, and *Myc* expression level is directly proportional to the selection signal strength (*3, 44–47*). Notably, we observed higher expression of *Myc* and mTORC1 in positively selected LZ/DZ intermediate cells (C4) from SMNP– and alum-pSer/SMNP-immunized mice compared to alum-pSer alone (**Fig. 2I**, **fig. S2G**); the target genes of Myc and mTORC1 (**Data S1**, retrieved from (*48, 49*) were also significantly upregulated in alum-pSer/SMNP and trended higher in SMNP (p_*_Myc_*_-target_=0.077 and p_*_mTORC1-target_*=0.058) compared to alum-pSer (**Fig. 2J, fig. S2G**). B cells that capture more antigens and receive stronger T cell help signals in the LZ can undergo more proliferative cycles in the DZ (*44–46*). Prolonged activation of E2F family transcription factors was reported to drive this “inertial” cell division in the absence of extrinsic signals (*45, 50*). We found higher expression of activating E2Fs (*E2f1, E2f2, E2f3*) among positively selected (C4) and S phase (C5) cells from the combination alum-pSer/SMNP group compared to alum-pSer alone (**Fig. 2K, fig. S2G**). Collectively, these results suggest that the combination adjuvant may have enabled antigen-binding GC B cells to capture more antigen, leading to greater positive selection signals and more cycles of cell division in the DZ.

### Combining alum-pSer slow antigen delivery with SMNP augments both clonal expansion and clonal diversity

We next turned to paired heavy/light chain scBCR-seq to determine how alum-pSer and SMNP adjuvants impacted the repertoire of antigen-binding B cells recruited to the GC. After rigorous quality control, we recovered full-length heavy chain sequences from 2,286 MD39-binding B cells, full-length light chain sequences from 5,931 MD39-binding B cells, and paired BCR sequences from 1,460 MD39-binding B cells; on average, we recovered BCR sequences from 7, 9, and 14 mice immunized with alum-pSer, SMNP, and alum-pSer/SMNP, respectively (**fig. S3A**). Recovered BCRs were distributed across phenotypic clusters (**fig. S3B**), and B cells from expanded clones were enriched in S-phase (**fig. S3C**). Greater proportions of GC B cells from SMNP and alum-pSer/SMNP immunized mice class switched to IgG isotypes, a finding consistent with ELISA analysis of serum Ig isotypes assessed at day 28 (**fig. S3D-E**).

A first striking observation was that clone sizes from mice immunized with the combination alum-pSer/SMNP vaccine were much larger than either individual adjuvant group, with twenty-seven clones comprised of 10 or more cells (**Fig. 3A**). By contrast, SMNP immunization elicited only six clones with more than 10 cells and alum-pSer primed very low levels of clonal expansion (**Fig. 3A**). As another measure of clonal expansion, we quantified clonal evenness for cells recovered from individual mice using Pielou’s evenness score (*J*) (*51*). This analysis revealed a significantly lower *J* score for the combination and SMNP compared to alum-pSer (**Fig. 3B**). Lower clonal evenness corroborates greater clonal expansion due to the expansion of a sub-portion of the overall clones (*52*). We next examined the number of unique clones in the GC and observed that substantially more clones were primed by the alum-pSer/SMNP compared to alum-pSer alone (9.8-fold more) or SMNP (2.5-fold more), and SMNP elicited 4.0-fold more clones than the alum-pSer group (**Fig. 3C**). Plotting recovered clonotypes from each mouse ranked by their clone sizes revealed that alum-pSer/SMNP combination vaccine simultaneously augmented clonal expansion of individual clones and recruited a greater quantity of unique clones into the GC reaction (**Fig. 3D**). Thus, the combination adjuvant immunization both recruited more clones to the GC and triggered greater clonal expansion.

**Fig. 3.**
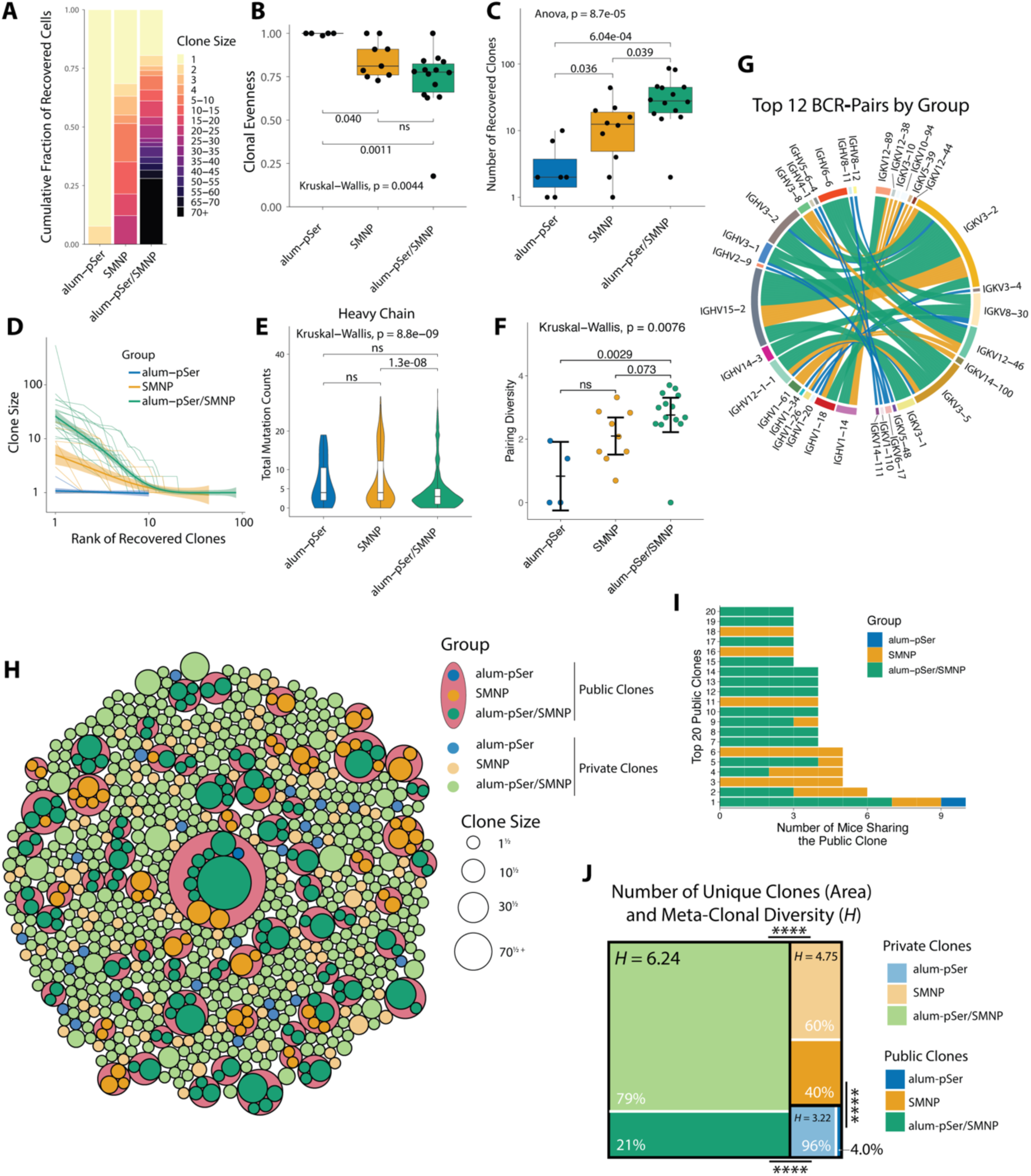
scBCR-seq analysis of MD39-binding germinal center B cells 14 days post-immunization. (**A**) Clone size distributions in each vaccine group. (**B**) The clonal evenness score computed by Pielou’s index. Only mice with recovered IgH sequences were included in the analysis. (**C**) Number of unique clones recovered per mouse across vaccine groups. Statistical significance was determined by two-way ANOVA followed by Tukey’s multiple comparisons test. (**D**) Clone recovery curves for each immunization mouse. Spline regression models were fit for each vaccine group. The shaded areas indicate 95% confidence intervals. (**E**) Heavy chain SHM counts. (**F**) Clonal heavy and light chain V gene pairing diversity per mouse, calculated by the Shannon diversity index. Shown are mean ± s.e.m. (**G**) Top 12 BCR pairs from each vaccine group. Each chord on the diagram represents one clone, and the colors indicate the vaccine groups. (**H**) Recovered meta-clonotypes. Private clones are illustrated as individual circles in lighter colors. Public clones are enclosed by coral-colored circles and illustrated in darker colors. The size of each circle is proportional to the square root of clone sizes. (**I**) Top 20 public clones. (**J**) Treemap illustration of meta-clonotypes. The areas of black-bordered boxes are proportional to the relative numbers of recovered meta-clonotypes per group. The areas of inner boxes are proportional to the percentage of private and public clones in each group. Meta-clonal diversity was labeled and computed by Shannon’s diversity index (*H*). Statistical significance for diversity indices was determined by Hutcheson’s t-test followed by Bonferroni multiple comparisons correction. **** p<0.0001. For (**B, E-F**), p values were computed with Kruskal-Wallis analysis of variance followed by Dunn’s post hoc test.

### Alum-pSer/SMNP vaccination enhances GC repertoire diversity

Many bnAbs for HIV have extensive SHM and require specific heavy and light chain V genes (*8–11, 13, 14*). Vaccines capable of recruiting diverse B cell clones would be advantageous to increase the likelihood of priming rare precursors capable of evolving toward broad neutralization (*10, 11, 13, 16, 18*). We thus sought to evaluate the BCR heavy and light chain SHM and pairing diversity.

We first counted synonymous and replacement nucleotide mutations in the recovered GC BCRs based on inferred germline sequences and found comparable heavy and light chain mutation counts across the vaccine groups (**Fig. 3E, fig. S3F**), consistent with prior work suggesting that degrees of SHM are influenced more by elapsed time post-immunization rather than vaccine formulation (*16, 17, 43*). While the SMNP group showed a statistically significant increase in heavy chain SHM counts over the alum-pSer/SMNP group, the effect size was one nucleotide.

To quantify heavy and light chain pairing diversity, we collapsed cells into their respective clones and aggregated clones by V gene pairs. We calculated the Shannon diversity index (*H*) for each group (*53*), where each unique V gene pair represents one species and the number of clones using the pair represents the abundance of that species. The calculation showed increasing diversity scores in the order of alum-pSer/SMNP > SMNP > alum-pSer (**fig. S3G-I**). The same calculation was done for each mouse, and the same hierarchy was observed, with the combination adjuvant showing significantly greater BCR pairing diversity than alum-pSer alone and a trend of greater diversity than SMNP alone (p=0.073, **Fig. 3F**). To determine whether this result reflected the larger size of GCs primed by the combination adjuvant or an emergent property due to the alum-pSer/SMNP formulation, we performed sub-sampling analyses on alum-pSer/SMNP (where 1222 paired BCR sequences and 303 unique clones were recovered, respectively) and SMNP (225 paired BCR sequences recovered, 104 clones) groups by random sampling either 200 BCRs or 100 clones from each group, calculating pairing diversity scores for each mouse, performing a two-tailed t-test, and repeating the process 10,000 times to generate distributions of p-values (**fig. S3J**). Neither analysis showed statistical significance, suggesting that the increasing trend of BCR pairing diversity was primarily correlated with overall GC size.

We observed overlaps among the most frequently used BCR pairs across the vaccine groups (e.g., Ighv3-2—Igkv3-2, Ighv12-1-1—Igkv3-5, and Ighv15-2—Igkv3-2, **Fig. 3G**), which motivated us to perform an analysis for public clones. Based on their use of (1) the same V and J genes, (2) the same HCDR3 length, and (3) a similarity threshold of their HCDR3 amino acid sequences, we grouped B cell clones into meta-clonotypes across mice and vaccine formulations. We defined “public clones” as meta-clonotypes encompassing clones from more than one mouse and the rest as “private clones” (**Fig. 3H-I**). This analysis revealed that 4%, 40%, and 21% clones from alum-pSer, SMNP, and alum-pSer/SMNP were grouped into public clones, respectively (**Fig. 3J**). Among the top 20 public clones, the largest public clone enclosed B cells from all three groups, and 4 public clones from SMNP and alum-pSer/SMNP (**Fig. 3I**). In total, thirty-nine public clones were observed uniquely from the alum-pSer/SMNP group, 2.1-times more than SMNP-unique public clones, suggesting that mice immunized with alum-pSer/SMNP developed more diverse BCR features which were positively selected for recognizing MD39 (**Fig. 3H**).

To gauge meta-clonal diversity, we calculated the Shannon diversity index (*H*) for each vaccine formulation, in which the abundance of each species (unique meta-clonotype) was the amount of enclosed private clones from each formulation (private clones were counted as meta-clonotypes with an abundance of 1). The alum-pSer/SMNP group exhibited the greatest meta-clonal diversity (*H* = 6.24), followed by SMNP (*H* = 4.75) and alum-pSer (*H* = 3.22) (**Fig. 3J**). Altogether, we found that mice vaccinated with alum-pSer/SMNP induced GCs with greater numbers of unique private and public clones. These results suggest that the repertoire of BCRs recruited to GCs following alum-pSer/SMNP immunization is more diverse.

### Combining alum-anchored immunogens with SMNP promotes trafficking of antigen to lymph node follicles

The scRNA-seq and scBCR-seq analyses showed that combining alum-pSer and SMNP adjuvants increased GC B cell clonal expansion, repertoire diversity, total number of B cell clones recruited to the GC, and increased the proportion of B cells cycling in the DZ– a surprising breadth of effects on the GC response. We thus next sought to investigate potential mechanisms underlying these effects of alum-pSer/SMNP immunization. The observation of S-phase enrichment in alum-pSer/SMNP GC B cells inspired a hypothesis based on our prior findings: heavily glycosylated antigens like HIV Env trimers, when displayed on the surface of nanoparticles, trigger complement deposition *in vivo* via the lectin pathway, resulting in complement-dependent trafficking of the particles to FDCs (*54, 55*). We hypothesized that alum particles bearing many pSer-anchored trimers could trigger a similar process for trafficking the antigen to FDCs. In parallel, we also previously showed that SMNP triggers rapid depletion of subcapsular sinus macrophages and increased antigen accumulation in dLNs (*25*). These two complementary mechanisms of action may be synergistic in shepherding more MD39 antigen into the follicles and onto the FDC network, which might explain the enrichment of S-phase GC B cells observed in the scRNAseq analysis, because greater antigen uptake is correlated with more cycles of cell division in the DZ (*44*) (**Fig. 2G-H, fig. S3C**).

Motivated by these ideas, we evaluated the localization of fluorescently labeled MD39 trimer in dLNs following vaccination with alum-pSer and SMNP. We chose to focus on comparing antigen trafficking of alum-pSer vs. alum-pSer/SMNP because we had previously shown that immunization with Env trimers and SMNP alone does not lead to significant antigen accumulation on FDCs in a primary immunization (*16, 56, 57*). As a control, we also assessed antigen biodistribution in LNs for SMNP and alum mixed with MD39-Ser_4_, a trimer conjugated with a non-phosphorylated serine tag that cannot undergo ligand exchange binding to the alum particles. Mice were immunized with these 3 different vaccines, and inguinal dLNs were isolated at varying time points for histological imaging. Following immunization with either alum/MD39-pSer_4_ or MD39-Ser_4_ combined with both alum and SMNP, minimal antigen accumulation on the FDC network (or at other locations in the LN) was observed at any examined time point (**Fig. 4A-B**). By contrast, when MD39 was bound to alum via a phosphoserine linker and combined with SMNP, substantial antigen accumulation was detected on LN FDCs beginning at day 14, and persisted through day 35 (**Fig. 4C**). Higher magnification imaging showed that this antigen signal colocalized with FDC dendrites (**Fig. 4D**). Quantification of the FDC-localized antigen signal from multiple follicles of multiple LNs over time showed that following alum-pSer/SMNP immunization, antigen accumulation rose sharply between day 7 and day 14, was maintained through day 21, and then slowly decayed thereafter (**Fig. 4E**). This analysis revealed that the combination adjuvant vaccination elicited at least 15-fold greater antigen accumulation on FDCs compared to immunization with alum/MD39-pSer_4_ alone. To investigate the mechanistic basis of antigen accumulation on FDCs following alum-pSer/SMNP immunization, we evaluated the localization of fluorescently labeled MD39 trimer in dLNs of mice lacking the C3 component of the complement system (C3 KO) (**Fig. 4F-G**). Strikingly, we observed a substantial reduction in antigen colocalization with FDCs in C3 KO mice compared to wild type C57BL/6 mice, suggesting that the trafficking of antigen to the FDCs following alum-pSer/SMNP immunization is complement-dependent.

**Fig. 4.**
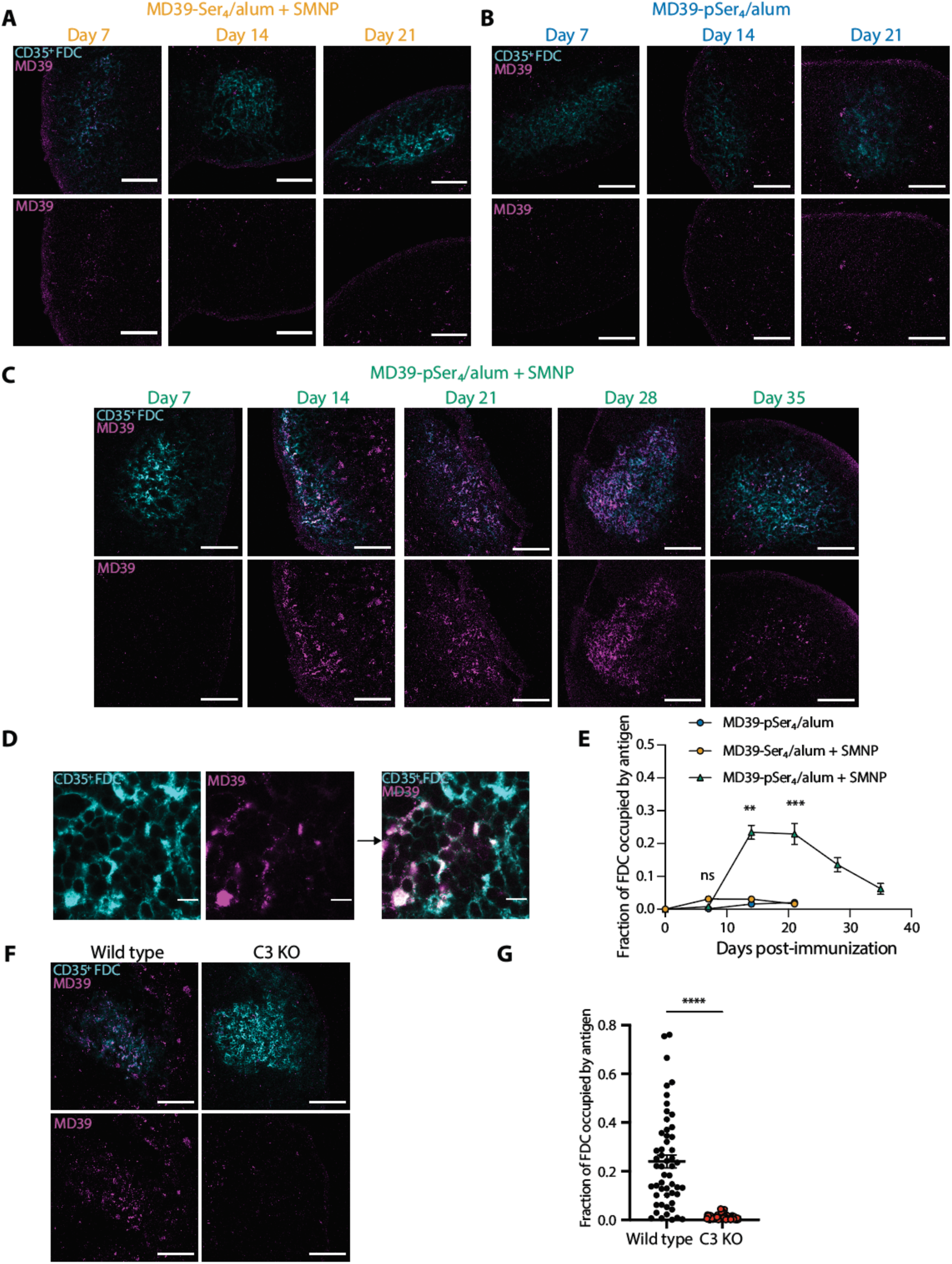
Env trimer antigen administered with alum anchoring and SMNP accumulates on the follicular dendritic cell network over time. BALB/c mice (*n*=3/group) were immunized with 10 μg labeled MD39-Ser_4_ (**A**) or MD39-pSer_4_ (**B**) combined with 100 µg alum, or MD39-pSer_4_ combined with 100 µg alum and 5 µg SMNP (**C**). Draining inguinal lymph nodes were isolated at the indicated timepoints, flash frozen, and cryo-sectioned. Shown are representative images of antigen on the lymph node CD35^+^ follicular dendritic cell (FDC) networks. Overlay of antigen and FDC shown in top row of images, with antigen signal alone shown in the bottom row of images. Scale bars, 100 µm. (**D**) Shown are representative 100x objective lens images of CD35^+^ FDC staining and MD39 antigen, with overlay on right. Scale bars, 10 µm. (**E**) The fraction of FDCs occupied by antigen was calculated for each timepoint and immunization. Values plotted are mean ± s.e.m. Statistical significance was determined by one-way ANOVA followed by Tukey’s multiple comparisons test. (**F**) Wild type C57BL/6 and C3 KO mice (*n*=3/group) were immunized with 10 μg labeled MD39-pSer_4_ combined with 100 µg alum and 5 µg SMNP. Draining inguinal lymph nodes were isolated 14 days after immunization, flash frozen, and cryo-sectioned. Shown are representative images of antigen on the lymph node CD35^+^ follicular dendritic cell (FDC) networks. Overlay of antigen and FDC shown in top row of images, with antigen signal alone shown in the bottom row of images. Scale bars, 100 µm. (**G**) Fraction of FDCs occupied by antigen for each mouse strain. Values plotted are mean ± s.e.m. Statistical significance was determined by Mann Whitney test. ns p>0.05, * p<0.05, ** p<0.01, *** p<0.001, **** p<0.0001.

### Env trimer antigen delivered to follicles via combined alum-pSer/SMNP vaccination is retained in a highly intact state on FDCs

Antigen localizing to the sinuses and interfollicular regions of the LN undergoes rapid degradation over the first few days post-immunization, while antigens captured on FDCs can be retained in a highly intact state over time due to spatially compartmentalized protease activity in LNs (*57*). To assess potential degradation of the MD39 trimer *in vivo* following combination alum-pSer/SMNP immunization, we applied a fluorescence resonance energy transfer (FRET)-based approach we previously developed to track antigen integrity in LNs (*57*). MD39 antigen was labeled with ∼6 total dyes per trimer (∼3 Cy3 donor and ∼3 Cy5 acceptor dyes) for FRET imaging. When the antigen undergoes proteolytic degradation, the donor and acceptor dyes become separated, leading to reduced FRET signals proportional to the degree of degradation (**fig. S4A**) (*57*).

We first confirmed that this degree of dye labeling did not significantly alter the alum binding behavior or antigenicity profile of the trimer (**fig. S4B-C**). We next tested whether the binding of dye-labeled MD39-pSer_4_ trimer to alum affected the measurement of intermolecular FRET, using an acceptor photobleaching approach to measure energy transfer. We imaged FRET dye-labeled free MD39 or alum-anchored MD39-pSer_4_ adsorbed on glass coverslips and observed an enhancement in donor emission following acceptor photobleaching indicative of FRET (**fig. S4D-E**). The histograms of FRET efficiencies measured pixel by pixel for free vs. alum-bound trimer overlapped, indicating no significant impact of alum binding on the FRET signal (**fig. S4F**). FRET was only observed when antigens were labeled with both Cy3 and Cy5 on the same trimer, and no intermolecular FRET was observed when Cy3-labeled MD39-pSer_4_ trimers were co-loaded with Cy5-labeled MD39-pSer_4_ on alum particles, or when alum particles loaded with Cy3-labeled MD39-pSer_4_ were mixed with alum particles carrying Cy5-labeled MD39-pSer_4_ (**fig. S4G**). These controls indicate that FRET detected *in vivo* should reflect intact trimeric antigen and not intermolecular FRET between adjacent trimers loaded on alum or bound to FDCs. In addition, the FRET efficiency was not influenced by adding Cy3/Cy5-specific antibodies to the labeled trimer (**fig. S4H**), suggesting that potential anti-dye antibody responses which could theoretically be elicited by dye-labeled MD39 immunization would not alter the FRET efficiency readout. As expected based on our previous work, a decline in FRET efficiency was observed following the incubation of FRET-labeled, alum-bound trimer with the promiscuous protease trypsin (**fig. S5A-B**). This correlated with a reduction in the binding of antibodies targeting the interface/fusion peptide, V3 epitopes, and CD4 binding site on the trimer as measured by ELISA (**fig. S5C**). Overall, these data indicated that FRET-based imaging is sensitive to detect changes in the structural integrity of the trimer.

Having established the ability of the FRET assay to track MD39 trimer integrity when bound to alum, we next evaluated trimer stability in the context of alum-pSer and SMNP immunizations *in vivo*. Given that alum-bound trimer is expected to slowly clear from the injection site over time, we first investigated antigen stability at the injection site. To address this, mice were immunized subcutaneously with FRET dye-labeled MD39-Ser_4_ or MD39-pSer_4_ mixed with alum and SMNP adjuvants, and injection site tissues were isolated at varying time points post-immunization. While only low levels of MD39-Ser_4_ remained at the injection site by day 7 (**Fig. 5A**), alum-anchored MD39-pSer_4_ was still detectable at high levels at this time point and remained detectable at the injection site through day 21 (**Fig. 5A**), consistent with prior whole animal fluorescence imaging studies (*20*). Acceptor photobleaching FRET was used to determine the fraction of intact antigen at the injection site for each condition and timepoint, revealing that while there was a decline in antigen stability over time, ∼50% of alum-anchored MD39-pSer_4_ was still intact at the injection site 21 days after immunization (**Fig. 5B-C**).

**Fig. 5.**
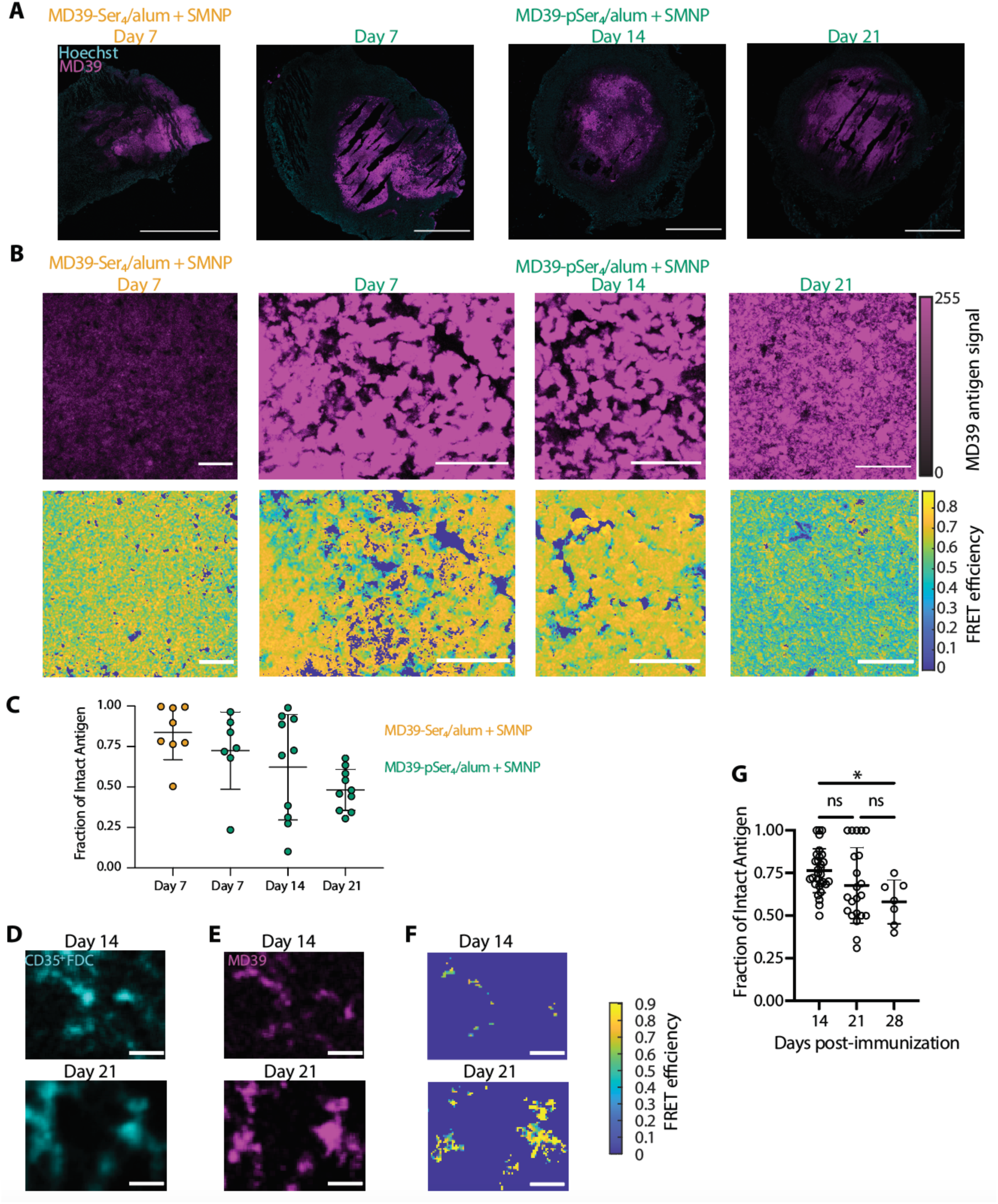
Env trimer antigen accumulated on FDCs following alum-pSer/SMNP immunization is highly intact. (**A**) BALB/c mice (*n*=3/group) were immunized with 10 μg of FRET dye-labeled MD39-Ser_4_ or MD39-pSer_4_ combined with 100 µg alum and 5 µg SMNP. Injection sites were isolated at the indicated timepoints, flash frozen, and cryo-sectioned. Shown are representative images of antigen and Hoechst nuclei staining at the injection site. Scale bars represent 1000 µm. (**B**) Representative pre-bleach acceptor images. These regions underwent acceptor photobleaching, enabling the calculation of FRET efficiencies, shown as a heatmap below. Scale bars, 50 µm. (**C**) The fraction of intact antigen at the injection site was calculated based on the antigen FRET efficiencies. Values plotted are mean ± s.d. Representative (**D**) CD35^+^ FDC and (**E**) pre-bleach acceptor images in the draining lymph node FDC following immunization with MD39-pSer_4_. Scale bars, 10 µm. These regions underwent acceptor photobleaching, enabling the calculation of FRET efficiencies, shown as a heatmap in (**F**). (**G**) The fraction of antigen that is intact in the follicles was calculated based on these FRET efficiencies. Values plotted are mean ± s.d. Statistical significance was determined by one-way ANOVA followed by Tukey’s multiple comparisons test. ns p>0.05, * p<0.05, ** p<0.01, *** p<0.001, **** p<0.0001.

In parallel, we imaged antigen accumulated on the FDC network in draining inguinal lymph nodes. Strikingly, a majority of antigen that accumulated on the FDCs following alum-pSer/SMNP immunization was non-degraded and remained intact through day 28 (**Fig. 5D-G**). Altogether, these data indicate that unlike alum-pSer or SMNP adjuvants individually, the combination alum-pSer/SMNP immunization leads to pronounced antigen targeting to FDCs over time, and this antigen is retained in a highly intact state for at least one month post-immunization, all factors that would be expected to help augment recruitment of trimer-specific B cells to the GC response.

## DISCUSSION

Slow-delivery vaccine approaches promote early immune complex formation and sustained antigen delivery to B cells, which has been shown to increase the diversity of recruited clones and correlated with enhanced neutralization breadth (*15–17*). On the other hand, particulate display of antigens is conducive to BCR crosslinking and B cell activation, which has been shown to facilitate the activation and recruitment of diverse precursors B cells, restrict access to base-proximal epitopes, and cultivate the maturation of low-affinity precursors (*54, 56, 58, 59*). These distinct (and we hypothesized, complementary) mechanisms of action underlying slow delivery and particulate antigen formulations inspired our initial studies examining the alum-pSer/SMNP adjuvant combination. Here, we showed that this combination adjuvant vaccine, which incorporates slow-delivery, multivalent anchoring of immunogens onto alum, and a potent adjuvant, enhanced multiple facets of the humoral immune response including greater GC B cell, Tfh cell, and serum IgG antibody responses. Although mice are generally unable to elicit bnAb-lineage B cell responses due to their short CDR3 domains (*60*), we found that this combination adjuvant strategy recruited a more clonally expanded and diverse B cell response and resulted in a repertoire with broader BCR characteristics to recognize the MD39 trimer immunogen—all features expected in humans to promote priming of rare precursor B cells that are needed for bnAb responses against HIV and other infectious diseases.

One important consideration for vaccine slow delivery approaches is the possibility of antigen degradation over time following administration, either at the injection site or en route to draining lymph nodes. Despite the promise of the alum-pSer approach to promote sustained delivery of antigen *in vivo*, when exposed to physiological conditions, including proteases and thermal stress for extended periods of time, antigen breakdown could occur that could influence B cell competition and immunodominance, diverting responses toward vaccine-irrelevant breakdown product epitopes (*61*). Previous studies have pointed to both positive and negative impacts of alum on antigen integrity and stability: adsorption of proteins onto alum has been reported to result in protein unfolding, potentially due to the nature of the protein-alum interactions or the duration of the binding (*62–64*), while others point to a stabilizing role for antigen adsorption on alum for antigen integrity, particularly when exposed to thermal stress (*65*). Extensive efforts have sought to stabilize HIV Env immunogens through molecular engineering (*66*), and we hypothesized that alum-anchoring of pSer-Env trimers might partially shield immunogens from protease-mediated degradation. Using a FRET imaging-based approach to track antigen integrity over time, we found that a majority of alum-bound pSer-trimer remained intact for at least 28 days post-injection (**Fig. 5G**). Since our approach utilizes a phosphoserine peptide tag to mediate antigen-alum ligand exchange interactions, it is possible this alum-pSer platform may be less susceptible to antigen-specific stability phenomena observed in the literature.

Another important facet of vaccine design is the ability of antigens to traffic to the FDC network. Previous studies have demonstrated that antigen particles decorated with complement (either due to innate immune recognition or immune complex formation (*67–69*)), efficiently accumulate on FDCs, leading to enhanced GC and serum antibody responses (*54, 56*). Antigens captured by FDCs shape the B cell response, as FDCs present antigens to B cells in the follicle where activated B cells undergo proliferation and SHM to generate high-affinity antibodies (*69, 70*). Notably, soluble HIV trimer immunogens delivered as bolus injections predominantly localize in interfollicular regions via SIGN-R1^+^ lymph node macrophages, which capture the trimer from the afferent lymph, rather than trafficking to FDCs (*54, 71*). Being able to deliver antigens onto the FDC network is of interest because FDCs can recycle and protect antigens captured on their dendrites (*67*) and the follicles are sanctuary sites with low protease activity in lymph nodes where antigens are protected from proteolytic degradation (*57*). It is known that protein nanoparticles or immune complexes decorated with complement are shuttled to FDCs in a complement– and complement receptor-dependent manner, mediated by noncognate B cells picking up the complement-decorated antigen and transferring it to FDCs (*54, 68, 72–74*). In the present case, there are at least 3 possible pathways for alum/pSer-trimer particles to trigger complement deposition: (i) early antibody responses elicited over the first 1-2 weeks could promote the formation of immune complexes as alum/pSer-trimer complexes slowly drain from the remaining injection site depot to the dLN, similar to effects observed with repeat-injection immunizations (*15, 16*); (ii) the alum particles may directly activate complement (*75, 76*) leading to C3 decoration of the alum/pSer-trimer complexes; or (iii) complement deposition could be triggered by mannose binding lectin recognition of trimer-decorated alum particles (*54*). While we show here FDC homing of antigen following the combination adjuvant vaccination is dependent on complement, further dissecting whether one or more of these pathways governs this response remains an open question for future work. SMNP is expected to amplify this antigen delivery process by causing early depletion of subcapsular sinus macrophages that limit antigen entry into the dLN.

In conclusion, these studies demonstrate how combining slow antigen delivery via immunogen anchoring on alum with a potent saponin/TLR agonist adjuvant can alter antigen biodistribution in lymph nodes, leading to a sustained buildup of intact antigen captured in B cell follicles. This alteration in antigen delivery correlates with substantial changes in the composition of the germinal center response triggered by the combination of these two adjuvants: including recruitment of a significantly more diverse set of B cell clones to the GC, which also undergo greater clonal expansion compared to vaccines using either adjuvant alone. More broadly, this work reinforces the idea that combining advancements in vaccine delivery with developments in adjuvant technologies can enable the induction of distinctly different immune responses to immunization via novel mechanisms of action.

## MATERIALS AND METHODS

### Study design

The objective of these studies was to investigate the mechanism by which the combination of alum-anchored pSer-modified HIV Env immunogens with SMNP adjuvant elicits significantly improved humoral immune responses, as a clinically translatable approach to promote slow delivery of vaccine following a single injection. Mice were immunized and subsequent humoral immune responses (germinal center responses, T follicular helper responses, and serum antibody responses) were assessed by flow cytometry, ELISA, and single-cell RNA sequencing. Antigen trafficking studies were completed using fluorescently labeled antigens and imaged by confocal microscopy using acceptor photobleaching to track antigen integrity via FRET. This acceptor photobleaching method avoids donor and acceptor crosstalk and is not influenced by dye concentrations or ratios. Group sizes were selected based on effect sizes seen in prior studies.

### Antigen production and pSer conjugation

MD39 immunogens with or without a free C-terminal cysteine and containing a positively charged, non-polyhistidine amino acid sequence (Lys-Lys-Lys) at the C-terminus of the trimer with or without a filled glycan hole at residues N241 and N289 (*20, 77, 78*) were synthesized as described previously (*27, 79*). Briefly, genes encoding MD39 HIV Env gp140 were cloned into pHLsec by Genscript and co-transfected with human furin in a pcDNA3.1 plasmid using a 2:1 trimer:furin DNA ratio with polyethylenimine into FreeStyle 293-F cells (ThermoFisher) and incubated for 6 days. The cultures were centrifuged and the supernatants containing MD39 were harvested and purified using a HisTrap HP column (Cytiva Life Sciences) with an AKTA FPLC system (Cytiva Life Sciences) for immunogens expressed with a polyhistidine linker and a 2G12 immunoaffinity column for MD39 immunogens without a polyhistidine linker. The immunogens were further purified by size-exclusion chromatography with an S200 Increase column (Cytiva Life Sciences) in TBS at flow rate of 0.5 mL/min. Size exclusion chromatography multi-angle light-scattering (SECMALS, DAWN HELEOS II and Optilab T-rEX Wyatt Technology) was then used to confirm the trimer molecular weights.

Immunogens expressed with a free terminal cysteine were reduced at 1 mg/mL with 10 molar equivalents of tris(2-carboxyethyl)phosphine (TCEP, ThermoFisher) in TBS and incubated at 25°C for 10 minutes. TCEP was subsequently removed from the reduced protein solution using Amicon Ultra Centrifugal Filters (10 kDa MWCO, Millipore Sigma) in tris-buffered saline (TBS, Sigma Aldrich), and 1 mg/mL reduced antigen was reacted with 5 molar equivalents of Ser_4_-maleimide or pSer_4_-maleimide linkers for 16 hours at 4°C in TBS (pH 7.2-7.4). Free peptide linker was subsequently removed using 10 kDa MWCO centrifugal filters in TBS, and pSer-antigen was buffer exchanged to PBS.

pSer_4_-conjugated cytochrome C used for antigenicity profiling of immunogens was prepared as described (*18*), using cytochrome C from Saccharomyces cerevisiae (Sigma Aldrich). The number of pSer residues conjugated to the antigen was assessed using the Malachite Green Phosphoprotein Phosphate Estimation Assay Kit (Thermo Scientific) against a standard curve of pSer-maleimide linker. Signal from pSer-antigen was compared to the background from an unconjugated antigen control.

### Animals and immunizations

Experiments and handling of mice were conducted under federal, state, and local guidelines under an Institutional Animal Care and Use Committee (IACUC)-approved protocol. Female 6-8-week-old BALB/c, C57BL/6, and C3 KO mice were purchased from the Jackson Laboratory (stock no. 000651, 000664, and 029661, respectively). Immunizations for sequencing studies were prepared by mixing 5 μg of antigen with a glycan hole at residues N241 and N289 and 50 μg of alum in 100 μL sterile tris-buffered saline (TBS, Sigma Aldrich) per mouse. Immunizations for histology and FRET studies were prepared by mixing 10 μg of antigen with a filled glycan hole at residues N241 and N289 and 100 μg of alum in 100 μL sterile tris-buffered saline (TBS, Sigma Aldrich) per mouse. Antigen was loaded onto alum for 30 minutes on a tube rotator prior to immunization. For alum-pSer/SMNP combination vaccines, antigen was first loaded onto alum for 30 minutes on a rotator, after which 5 μg of SMNP was added and incubated with antigen-alum formulations for 30 minutes prior to immunization. This dose of SMNP corresponds to 5 µg of Quil-A saponin and 0.5 µg MPLA. For sequencing studies, BALB/c mice were immunized subcutaneously at the tail base with 50 μL on each side of the tail base with one of three formulations: alum-pSer (5 µg MD39-pSer_4_ and 50 µg alum total), SMNP (5 µg MD39 mixed with 5 µg SMNP) or alum-pSer/SMNP (5 µg MD39-pSer_4_ combined with 50 µg alum and 5 µg SMNP).

### scRNA-seq study design, processing, and analysis

The inguinal LNs of vaccinated mice (n=14 animals/group) were isolated 14 days post-immunization. LNs were mashed and passed through a 70um filter to obtain single-cell suspensions. Cells were stained for viability (ThermoFisher Live/Dead FixableAqua) and with antibodies against CD3e (BV711, BioLegend, 145-2C11clone), CD19 (APC, BioLegend, 6D5 clone), B220 (PE-Cy7, BioLegend RA3-6B2 clone), CD38 (FITC, BioLegend 90 clone), GL7 (PerCP-Cy5.5, BioLegend GL7 clone), and labeled with TotalSeq-B cell hashing antibodies (BioLegend). Antigen-specific staining was completed using biotinylated MD39 conjugated to streptavidin-BV421 (BioLegend) and streptavidin-PE (BioLegend). Antigen double-positive GC B cells were sorted on a BD FACS Aria (BD Biosciences) cell sorter and processed immediately following the SeqWell protocol (**Table S1**) (*28, 29*). scRNA-seq libraries were sequenced by Illumina NovaSeq and aligned to the mm10 reference genome using the STARsolo pipeline (version 2.4.0) (*80*). The gene expression count matrix and cell hashing sequence reads were processed and analyzed using Seurat (v4.1.0), CITE-seq-Count v1.4.2, Scanpy, and scVelo (*81–84*) (for detailed procedures, see Supplementary Materials and Methods).

Myc– and mTORC1-target genes (**Data S1**) were retrieved from the literature (*45, 48, 49*) and their gene set module scores were calculated using the AddModuleScore() function in Seurat (*81*). Data from Mathew et al., King et al., and Holmes et al. were obtained from ArrayExpress with accession numbers E-MTAB-9478, E-MTAB-9005, and from Gene Expression Omnibus database GSE139891, respectively (*41–43*). Differentially expressed genes of relevant clusters from these studies were mapped onto our scRNA-seq data using the AddModuleScore() function (*81*).

### scBCR-seq library generation, sequencing, processing, and analysis

The scBCR-seq library was prepared as described previously (*30*). In brief, the immunoglobulin (Ig) transcripts were enriched and amplified from the 3’-barcoded cDNA (**Table S2**). A set of V-gene primers (modified from Tillers et al. (*85*)) (**Table S3**), were used to further enrich immunoglobulin transcripts before sequencing (**Table S4-S5**). BCR sequence reads were processed with pRESTO (v0.5.13), Change-O (v0.4.6), UMI-Tools, and IgBlast (v1.14.0) (*86–89*) to reconstruct and annotate full-length BCR sequences that match corresponding single-cell transcriptomes (for detailed procedures, see Supplementary Materials and Methods).

Clonally related sequences were identified using the DefineClones.py function (Change-O v0.4.6) with a 90% CDR3 nucleotide similarity threshold determined by distToNearst() function from the Shazam package (*87*). Germline sequences were inferred using the CreateGermline.py function (Change-O v0.4.6) (*87*). Meta-clonotypes were identified based on the shared V gene, J gene, CDR3 junction length, and 94% CDR3 amino acid similarity threshold determined by distToNearst() function. Clone sizes, or the number of cells per clone, were calculated using the countClones() function (alakazam v1.2.1). Clonal evenness was quantified for each mouse by Pielou’s evenness index (*J*) (*51*). It was calculated as the Shannon Diversity Index (*H*) divided by the natural log of the total number of clones in each mouse (*53*). A *J* value of 1 means that all the clones in the mouse have the same number of cells.

SHM counts were aggregated using the observedMutations() (Shazam v1.1.2) function (*87*). The Circos plots were generated using the “circlize” package (*90*). BCR pairing diversity was calculated in the following steps: first, cells were collapsed into their respective clones. If a clone matched to more than one light chain V gene, the light chain V gene with the most cells in the clone was designated for the clone. Second, clones were aggregated into unique BCR pairs and the number of clones using a unique BCR pair was counted. Lastly, Shannon’s Diversity Index was calculated for each group, whereby each unique V gene pair was the “species” and the number of clones using the pair was the “abundance” for the “species”. The same calculation was also performed at the individual mouse level.

### Confocal microscopy

Imaging was performed on a Leica SP8 confocal microscope using white light and argon lasers with spectral emission filters with a minimum bandwidth of 10 nm. A 25X water-immersion objective was used unless otherwise indicated. Laser and detector settings were kept constant across all imaging timepoints for each antigen. FRET efficiency was calculated as previously described (*57*). Images were processed and analyzed in ImageJ (version 2.1.0/1.53c).

### Statistical Analysis

For sequencing analysis, statistical analysis was performed in the R software version 4.3.0. The specific statistical tests are indicated in the figure legends. All other data were plotted and all statistical analyses were performed using GraphPad Prism 9 software (La Jolla, CA). All graphs display mean values, and the error bars represent the standard deviation unless otherwise specified. No samples or animals were excluded from the analyses. Statistical comparison was performed using a one-way ANOVA followed by Tukey’s post-hoc test for single timepoint data with three or more groups and two-way ANOVA followed by Tukey’s post-hoc test for multi-timepoint longitudinal data unless otherwise indicated. Data were considered statistically significant if the p-value was less than 0.05.

### List of Supplementary Materials

Supplementary Materials and Methods

Fig S1 to S5

Data S1

Tables S1 to S5

## Acknowledgments

We thank the Koch Institute Robert A. Swanson (1969) BioMicro Center and Biotechnology Center for technical support, specifically the Histology, Flow Cytometry, and Microscopy cores. Select figures created with BioRender.com. DJI is an investigator of the Howard Hughes Medical Institute.

## Funding

National Institutes of Health grant P30-CA14051 (Koch Institute Core grant)

National Institutes of Health grant UM1AI144462 (ABW, WRS, DJI)

National Institutes of Health grant AI161818 (DJI)

National Institutes of Health grant AI161297 (DJI)

National Institutes of Health grant AI125068 (DJI)

National Institutes of Health grant P01AI048240 (DJI)

Ragon Institute of MGH, MIT, and Harvard (DJI)

Howard Hughes Medical Institute (DJI)

National Institutes of Health Fellowship F32 AI164829 (PY)

## Author contributions

KAR, YJZ, AA, DM, JCL, and DJI conceived the study plan. KAR synthesized the pSer linkers and SMNP. WRS designed the antigen constructs. KAR completed the in vitro characterization of the antigen constructs. YJZ and DM completed cell sorting, sequencing, and sequencing analysis. KAR, AA, LM, PY, JRG, PA, and MB completed the mouse studies. GG and GO completed the TEM imaging. KAR and AA completed fluorescence imaging. KAR, YJZ, and DJI wrote the manuscript. All authors reviewed and edited the manuscript. ABW, WRS, JCP, and DJI supervised the study.

## Data and materials availability

All data are available in the main text or the supplementary materials.

