## Supplementary Methods Figures S1-S5 and Tables S1-S5 for "Vaccines combining slow delivery and follicle targeting of antigens increase germinal center B cell clonal diversity and clonal expansion"

**This PDF file includes:**

Supplementary Materials and Methods
Figs. S1 to S5
Data S1
Tables S1 to S5

### Supplementary Materials and Methods

Phosphoserine peptide synthesis

$pSer_4$-maleimide and $Ser_4$-maleimide peptide linkers were synthesized using solid phase synthesis on low-loading TentaGel Rink Amide resin (0.2 meq/g, Peptides International) as described previously. Briefly, resin was deprotected with 20% piperidine (Sigma Aldrich) in dimethylformamide (DMF, Sigma Aldrich), and peptide couplings were performed with 4 equivalents of Fmoc-Ser(PO(OBzl)OH)-OH (Millipore Sigma) (or Fmoc-Ser(tBu)-OH for Ser linkers) and 3.95 equivalents of hexafluorophosphate azabenzotriazole tetramethyl uranium (HATU) for 2 hours at 25°C in dichloromethane and dimethyl formamide (DMF) (1:2 vol:vol). pSer residues were deprotected with 5% 1,8-diazabicyclo[5.4.0]undec-7-ene (DBU) in DMF for 45 minutes. Double couplings were performed for the third and fourth residues. An Fmoc-protected 6-unit oligoethylene glycol linker (Peptides International DPG-5750) was then coupled to the peptide, deprotected, and reacted with N-maleoyl-β-alanine (Sigma Aldrich). Completion of each deprotection and coupling step was confirmed by a ninhydrin test (Sigma Aldrich). pSer side chains were deprotected and the peptide was cleaved from the resin in 95% trifluoroacetic acid (Sigma Aldrich), 2.5% $H_20$, and 2.5% triisopropylsilane (Sigma Aldrich), for 2.5 hours at 25°C. The product was precipitated in 4°C diethyl ether (Sigma Aldrich) and dried under $N_2$, then purified by HPLC on a C18 column (Agilent Zorbax 300SB-C18) using 0.1 M triethylammonium acetate buffer (Glen Research) in an acetonitrile gradient. The peptide mass was confirmed by matrix-assisted laser desorption/ionization-time of flight mass spectrometry.

SMNP adjuvant synthesis

Saponin-MPLA nanoparticle (SMNP) adjuvant was prepared as previously described. Briefly, solutions at 20 mg/mL were prepared of cholesterol, DPPC, and PHAD MPLA (Avanti Polar Lipids) in 20% MEGA-10 detergent (Sigma). Quil-A saponin (InvivoGen) was dissolved in Milli-Q water at a final concentration of 100 mg/mL. These stock solutions were mixed at a mass ratio of 10:2:1:1 Quil-A:chol:DPPC:MPLA and diluted in PBS to a final cholesterol concentration of 1 mg/mL. The solution was equilibrated overnight at 25°C and then dialyzed for 5 days against PBS (5 L PBS volume for 50mg Quil-A scale; PBS replaced every 12 hours) using a 10kDa MWCO cassette (ThermoFisher) to remove detergent. The adjuvant was then sterile filtered and purified by FPLC using a Sephacryl S-500 HR size exclusion column (Cytiva Life Sciences). SMNP was concentrated to ~3 mg/mL relative to Quil-A using Amicon Ultra Centrifugal Filters (50 kDa MWCO, Millipore Sigma), and the final concentration of SMNP was determined using a cholesterol quantification assay (Sigma Aldrich).

Germinal center response analysis

Inguinal lymph nodes were collected from immunized mice at indicated time points. For germinal center analysis, cells were stained for viability (ThermoFisher Live/Dead Fixable Aqua) and against CD3e (BV711, BioLegend, 145-2C11 clone), B220 (PE-Cy7, BioLegend RA3-6B2 clone), CD38 (FITC, BioLegend 90 clone), and GL7 (PerCP-Cy5.5, BioLegend GL7 clone), with antigen-specific staining completed using biotinylated MD39 conjugated to streptavidin-BV421 (BioLegend) and streptavidin-PE (BioLegend). Samples were analyzed by flow cytometry on a BD Celesta and analyzed on FlowJo.

Serum ELISA

Serum was collected from mice retro-orbitally using capillary tubes and stored at -20°C until analysis. To determine serum IgG titers, Nunc Maxisorp plates (Invitrogen) were coated with Galanthus nivalis lectin (Sigma-Aldrich) at 2 µg/mL for 4 hours at 25°C and blocked with 2% BSA in PBS for 16 hours at 4°C. Plates were subsequently washed with 0.05% Tween-20 in PBS and incubated with 2 µg/mL unmodified or pSer-conjugated MD39 in 2% BSA in PBS for 2 hours at 25°C. Serum dilutions (1:10 dilution followed by 1:50 dilution with 1:4 serial dilutions) were incubated in the plate for 2 hours. Plates are washed again, incubated with a goat anti-mouse IgG HRP-conjugated secondary (BioRad) at 1:5000 dilution, and then developed with 3,3′,5,5′-tetramethytlbenzidine (ThermoFisher), stopped with 2N sulfuric acid, and immediately read (450nm with 540nm reference) on a BioTek Synergy2 plate reader. Isotype ELISAs followed the same protocol but used goat anti-mouse IgG1 HRP cross-adsorbed secondary antibody, goat anti-mouse IgG2a HRP cross-adsorbed secondary antibody, or goat anti-mouse IgG2b cross-adsorbed secondary antibody (Invitrogen) at 1:2000 dilution.

scRNA-seq sequencing, alignment, processing, and analysis

cDNA was sequenced by Illumina NovaSeq 6000 SP. Sequencing reads were demultiplexed and aligned to the mm10 reference genome using the STARsolo pipeline (version 2.4.0) *(74)* on Terra.Bio with default parameters except star_version = "2.7.10b", soloType = "CB_UMI_Simple", soloCBstart = 1, soloCBlen = 12, soloUMIstart = 13, soloUMIlen = 8, soloCellFilter= "TopCells 10000" for Seq-Well-specific libraries.

The gene expression count matrix was processed using the Seurat (v4.1.0) package in R *(75)*. The initial quality control filtered out genes that were detected in less than 3 cells and removed cells with less than 300 genes or greater than 10% mitochondrial genes. Cell hashing sequence reads were aligned to hashtag oligo (HTO) barcodes using CITE-seq-Count v1.4.2 *(76)*. The HTO count matrix was added to the Seurat object and normalized. The HTODemux() function was used to assign HTO to each cell. Only singlet cells by HTO assignment were kept for downstream analysis.

Cells were normalized using the NormalizeData() function. The cell cycle was predicted using the CellCycleScoring() function, and variable genes were identified using the FindVariableFeatures() function. The ScaleData() function was used to regress out RNA feature counts and percent of mitochondrial genes before performing principal component analysis (PCA) using the RunPCA() function. Fifteen principal components (PCs) and 500 decision trees were used for constructing the nearest-neighbor graph with the FindNeighbors() function. Forty neighboring points and fifteen PCs were used to generate uniform manifold approximation and projection (UMAP) with the RunUMAP() function. Unsupervised clustering was determined using Louvain clustering as implemented in the FindClusters() function. Differential gene expression analysis was performed using FindMarkers() and FindAllMarkers() functions with the Wilcoxon Rank Sum test and the p-values were adjusted using Bonferroni correction. A cluster of *Ighd*$^{+}$ *Zfp318*$^{+}$ *Sell*$^{+}$ *Klf2*$^{+}$ *Cmah*$^{+}$ GC cells was discarded as likely contaminating naïve follicular B cells *(35)*.

Exonic (spliced) and intronic (unspliced) transcripts were counted with STARsolo *(74)*, integrated with scRNA-seq counts, and normalized using the Scanpy package *(77)*. The dynamical model from the scVelo *(78)* package was used to estimate RNA velocity, where 30 PCs and 30 neighbors were used to calculate the moments for velocity estimation. The estimated velocity was visualized as vector fields projected onto the UMAP embeddings. Velocity-based pseudo-temporal relationships among cells were approximated using the scvelo.tl.latent_time() function to infer the real-time experienced by cells.

Enrichment of immunoglobulin transcripts

The enrichment of immunoglobulin (Ig) transcripts from Seq-Well 3'-barcoded whole transcript amplification (WTA) products was performed using xGen Lockdown reagents (IDT, cat. no. 1072281). Biotinylated probes for IGHM, IGHD, IGHG12, IGHG3, IGHA, IGKC, IGLC, and IGLC2&3 were synthesized by IDT and were used at a concentration of 1.5 uM (each; 7.5 uM for heavy chain and 4.5 uM for light chain in total) (**Table S2**). We performed the enrichment for heavy chains and light chains separately. 3.5 uL of Seq-Well WTA product was combined with 8.5 uL 2x hybridization buffer, 2.7 uL hybridization buffer enhancer, 0.8 uL Seq-Well WTA primer (40 uM), 1uL probe mix, and 0.5 uL mouse cot-1 DNA (Invitrogen, cat. No. 18440-016) were combined and let sit at room temperature for 5 minutes. The mixture was then incubated at 95C for 10 minutes and 65C for one hour. The pull-down steps follow exactly according to the remainder of the xGen lockdown protocol. 50 uL of Dynabeads M-270 Streptavidin (Invitrogen, cat. no. 65306) were used for each sample. At the end of this protocol, Ig-bound beads were mixed in 20 uL of water.

The enriched products were then amplified with PCR. Five PCR reactions for each enriched sample were performed with the following composition per reaction: 12.5 uL of 2x Kapa Hifi Hotstart Readymix (Roche, cat. no. KK2602), 8.5 uL water, 2.0 uL Seq-Well WTA primer (10 uM), and 2.0 uL of Ig-bound beads (**Table S1-S2**). The following PCR cycling conditions were used: 1 cycle of 95C, for 3 minutes; 25 cycles of 98C for 40s, 67C for 20s, 72C for 1 min; 1 cycle of 72C for 5 min. The reactions were then pooled and purified using a homemade SPRI reagent at a ratio of 0.80x for the light chain and 0.65x for the heavy chain.

scBCR-seq library generation and sequencing

We designed Nextera-IGHV, Nextera-IGKV, and Nextera-IGLV primer sets for mice based on Tiller et al *(79)* with modifications (**Table S3**). Reaction mixtures for heavy and light chains were composed of 12.5 uL 2x Kapa Hifi Hotstart PCR Readymix, 6.0 uL water, 2.5 uL primer mix (equimolar pooling of primers with a final concentration of 10 uM), and 4.0 uL of enriched product. Primer extension was performed with the following thermal program: 98C, 5 minutes, 55C, 30s, 72C, 2 min for light chains; 98C, 5 minutes, 60C, 30s, 72C, 2 min for heavy chains. The products were cleaned with SPRI at a ratio of 0.8x (light chain) or 0.65x (heavy chain) and were eluted into 12 uL of water.

Four reactions of library index PCR were performed per sample. Reactions were composed of 0.5 uL P5_index_TSO primer (10 uM), 0.5 uL P7_index_Nextera primer (10 uM), 12.5 uL 2x Kapa Hifi Hotstart Readymix, 9 uL of water, and 2.5 uL of primer extension product (**Table S4**). Amplification used the following cycling conditions: 1 cycle, 95C, 2 min; 14-20 cycles of 95C, 30s, 60C, 30s, 72C, 1.5 min; 1 cycle of 72C, 5 min. Reactions were pooled and purified using SPRI at a ratio of 0.8x for light chains and 0.65x for heavy chains.

Final products were assessed using an Agilent Tapestation and D5000 ScreenTape (Agilent, cat. #5067-5588). Light chain libraries usually show a clean peak around 1000bp, and heavy chain libraries usually show a clean peak around 1500bp. Libraries were sequenced on an Illumina MiSeq using the 600-cycle kits or on an Illumina NovaSeq SP using the 500-cycle kits (**Table S5**). First, the Seq-Well R1 primer was used to sequence the cell barcode and UMI (20 nt on MiSeq, 26 nt on NovaSeq). Then, custom sequencing primers specific for the BCR constant region were used to sequence the BCR using the index 1 read (300 nt on MiSeq, 252 nt on NovaSeq). The 8-nucleotide i5 index barcode was sequenced using the Seq-Well R2 index primer (NovaSeq) or no custom primer (MiSeq). Lastly, the Nextera R2 primer was used to sequence the remainder of the BCR with read 2 (300 nt on MiSeq, 252 nt on NovaSeq).

scBCR-seq library processing

The molecular identity (Read 1) and paired-end reads (Index Read 1 and Read 2) were assembled *in silico* with pRESTO (v0.5.13) and Change-O (v0.4.6) pipelines *(80, 81)* to reconstruct full-length BCR sequences that match corresponding single-cell transcriptomes. Reads with an average quality (Q) score below 21 were removed using the FilterSeq.py function. Cell barcode and UMI reads (Read 1) with a Hamming distance of up to one nucleotide mismatch were collapsed to correct potential sequencing errors using UMI-Tools *(82)*. The MaskPrimers.py function was used to annotate and filter reads with the correct isotype (Index Read 1) and V-region primer (Read 2); the corresponding primer sequences on the reads were masked as well. Only read pairs (Index Read 1 and Read 2) that passed both the FilterSeq.py and MaskPrimers.py processing steps were retained using the PairSeq.py function.

To alleviate library preparation-related technical errors or sequencing-related errors such as barcode swapping and amplicon truncation, the ClusterSeqs.py function was used to sub-cluster Index Read 1 (3'-isotype) and Read 2 (5'-V(D)J) reads separately for each unique Ig molecule (cell barcode + UMI). The cluster fraction was calculated as the number of reads in each sub-cluster divided by the total number of detected reads for the Ig molecule (cell barcode + UMI). Only the cluster of reads with a cluster fraction > 0.5 was kept. Next, the BuildConsensus.py function was used to collapse and call consensus sequences on Index Read 1 and Read 2 separately for each unique Ig molecule (cell barcode and UMI). The AssemblePairs.py function was used to assemble these paired-end consensus sequences into a single overlapping sequence. The resulting sequences were annotated with the AssignGenes.py function and then analyzed with IgBlast (v1.14.0) *(83)* using reference sequences provided by IMGT.

Ig molecular consensus sequences with fewer than six reads and more than four ambiguous "N" characters were discarded. BCR sequences were matched to single cells using the 12 bp single-cell barcodes. To define a consensus BCR sequence for each cell, we performed single-linkage clustering on the recovered IMGT-gapped sequences using Levenstein distance for each cell and clustered sequences less than five distances away. We extrapolate the cellular consensus sequence from the largest cluster by a recursive string consensus-building algorithm. If the resulting consensus retains ambiguous ("N") characters, we discarded the sequence with the fewest number of reads in the cluster and re-attempted to determine a consensus sequence until no ambiguous character remained. This process allowed us to combine information from multiple BCR molecules recovered from the same cell into a single cell-level consensus sequence.

Electron microscopy

100 μg of BG505 SOSIP, recombinantly expressed and purified by 2G12 affinity and size-exclusion chromatography as described previously *(85)*, was incubated with 375 μg SMNP for 1 hour at room temperature. After 1 hour, the mixture was diluted using Tris-buffered saline to ~0.03 mg/mL BG505 SOSIP concentration and a 3 μL drop was applied to carbon-coated copper mesh grids, blotted, and stained for 60 s with 2% (w/v) uranyl formate.

Automated data collection was set up using Leginon *(86)* on a 120 keV FEI Tecnai TF20 with a TVIPS TemCam F416 CMOS camera (62,000X magnification resulting in a 1.677 Å pixel size). Micrographs were saved and viewed in the Appion database *(87)*.

Antigen labeling

Cy3- and Cy5-labeled proteins were prepared as previously described *(57)* by diluting 1 mg/mL antigen solutions in PBS at 1:1 (v/v) in 0.2 M sodium bicarbonate buffer (Sigma Aldrich, pH 8.4). Stock solutions of Sulfo-Cyanine 3 and Sulfo-Cyanine 5 NHS esters (Lumiprobe) were prepared fresh in 0.2 M sodium bicarbonate pH 8.4 and added to the antigen. After reacting for 16 hours at 4°C, the solutions were desalted using Zeba Spin Desalting columns (40kDa MWCO, ThermoFisher), equilibrated in PBS, and 0.22 µm sterile filtered (Millipore Sigma). Antigens were stored at 4°C. The degree of labeling was determined by UV-vis spectroscopy using the extinction coefficients of the Sulfo-Cy3 and Sulfo-Cy5 NHS ester dyes, 162000 and 271000 $M^{-1}$ $cm^{-1}$, respectively.

Antigen-alum binding and release

Alexa dye-labeled antigen was mixed with Alhydrogel (alum, InvivoGen) in TBS at a 1:10 antigen:alum mass ratio at an alum concentration of 0.1 mg/mL, unless otherwise specified, for 30 minutes on a tube rotator at 25°C. To assess antigen binding to alum, samples were immediately centrifuged at 10,000x*g* for 10 minutes to pellet alum, and the fluorescence of the supernatant was measured against a standard curve of labeled antigen. To assess the release of antigen from alum, mouse serum was added to antigen-alum solutions post-loading to a final mouse serum concentration of 10 vol% and incubated at 37°C for 24 hours, unless otherwise specified. Samples were subsequently centrifuged at 10,000x*g* for 10 minutes to pellet alum, and the fraction of protein bound to alum was measured by fluorescence analysis of the supernatant using a Tecan Infinite M200 Pro plate reader.

Antigenicity profiling of immunogens

Antigenicity profiling of antigens was carried out by coating free trimer or alum-bound trimer on ELISA plates and assessing binding of serial dilutions of selected structure-sensitive monoclonal antibodies to the immunogens by ELISA. To capture alum on Nunc Maxisorp ELISA plates (Invitrogen), plates were first coated with $pSer_4$-conjugated cytochrome C at 2 µg/mL in PBS for 4 hours at 25°C. Alum was then added at 200 µg/mL in PBS to be captured by $pSer_4$-cytochrome C overnight at 4°C. To capture "free" MD39, plates were coated with mouse VRC01 antibody, rabbit 12N antibody, or Galanthus nivalis lectin (Sigma-Aldrich) at 2 µg/mL for 4 hours at 25°C and blocked with 2% BSA in PBS for 16 hours at 4°C. For both alum-coated and antibody-coated plates, plates were subsequently washed with 0.05% Tween-20 in PBS and incubated with 2 µg/mL unmodified or pSer-conjugated MD39 in 2% BSA in PBS for 2 hours at 25°C. Indicated monoclonal antibodies were added to antigen-coated plates at 5 µg/mL with 1:4 serial dilutions for 2 hours at 25°C. Plates were washed and antibody binding was detected with a goat anti-human HRP conjugated secondary antibody with minimal cross-reactivity (Jackson ImmunoResearch) at 1:5000 dilution in PBS containing 2% BSA and then developed with 3,3′,5,5′-tetramethylbenzidine (ThermoFisher), stopped with 2N sulfuric acid and immediately read (450nm with 540nm reference) on a BioTek Synergy2 plate reader. For antigenicity profiling after digestion with trypsin, plates were coated with rabbit 12N antibody and antigens were captured as described, followed by incubation with 0.1 mg/mL of TPCK-treated trypsin (ThermoFisher) at 37°C for 2 hours prior to antigenicity profiling ELISA.

*In vitro* trypsin digestion of antigen

Stability of FRET dye-labeled MD39 trimer was assessed *in vitro* using TPCK-treated trypsin (ThermoFisher). Trypsin was reconstituted at 2 mg/mL in 0.1 M ammonium bicarbonate buffer (Sigma Aldrich) pH 8 and subsequently mixed 1:1 (v/v) with 40 µg/mL of antigen in 0.1 M ammonium bicarbonate buffer pH 8. Samples were incubated at 37°C on a tube rotator for 2-16 hours. Antigens were subsequently coated on glass coverslips at 10 µg/mL for 16 hours at 4°C and washed in PBS prior to FRET imaging.

Tissue sectioning and staining

Inguinal lymph nodes were collected from immunized mice at varying timepoints post-immunization and placed into cryomolds containing optimal cutting temperature compound (Fisher Scientific) as previously described *(57)*. The cryomolds were subsequently flash frozen in 2-methylbutane (Millipore Sigma) pre-chilled in liquid nitrogen. Tissues were subsequently cryo-sectioned on a Leica CM1950 at 10 µm thickness and adhered to Superfrost Plus microscope slides (Fisher Scientific). Sections were stored at -80°C until use, at which point they were quickly thawed, fixed in 10% neutral buffered formalin for 8.5 minutes at 25°C, and washed for 10 minutes in PBS three times. Slides were subsequently incubated in blocking buffer (2% BSA with 0.1% Triton X-100 in PBS) for 30 minutes at 25°C and then stained. Injection sites were stained with CellMask Green (ThermoFisher) at 1:5000 for 25 minutes at 25°C, washed for 10 minutes in PBS three times followed by Hoechst at 1:10000 (ThermoFisher) for 10 minutes at 25°C. Lymph nodes were stained with BV421 anti-CD35 (BD Biosciences) at a 1:75 dilution for 2 hours at 25°C in blocking buffer. Slides were then washed for 10 minutes in PBS three times.PBS (9 µl) was added directly to the tissues before a #1.5 18 x 18 mm coverslip was placed over the sample and sealed with CoverGrip sealant (Biotium). Slides were stored at 4°C for a maximum of 1 week for imaging.

### Supplementary Figures

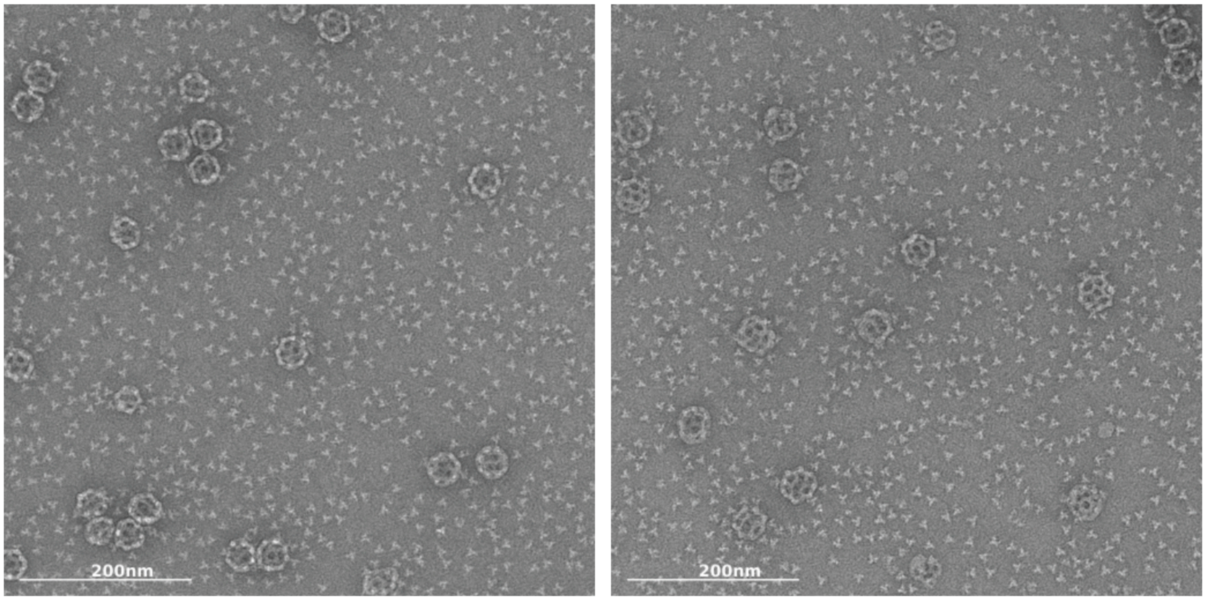

**Fig. S1. SMNP does not interact with HIV Env trimer.** Representative micrographs of 375 µg SMNP mixed with 100 µg BG505 SOSIP. Stained (with UF) and imaged after 1 hour incubation at 25°C. Scale bars represent 200 nm.

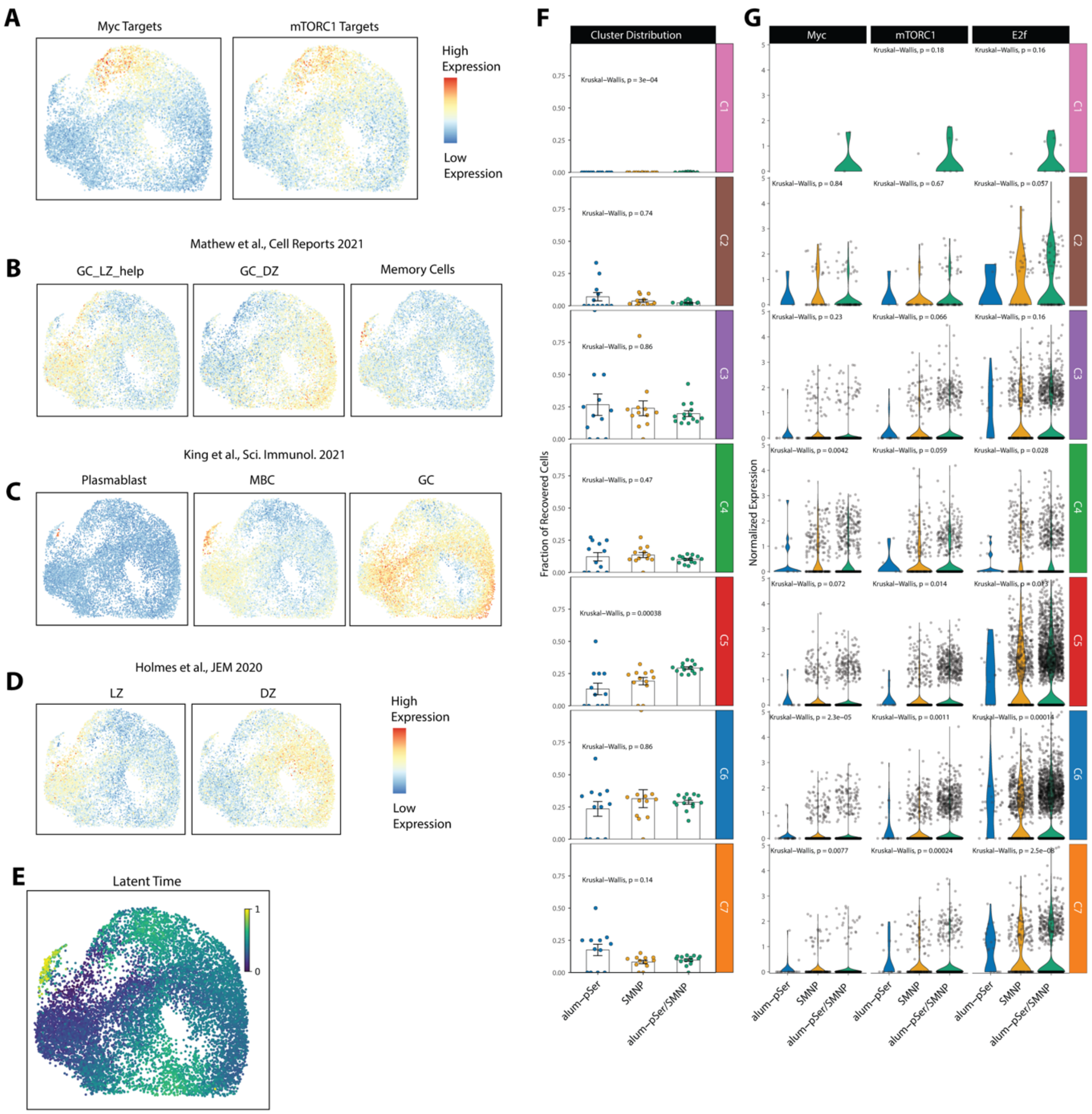

**Fig. S2. scRNA-seq analysis of MD39-binding GC B cells.** (**A**) UMAP projection of modules scores of *Myc*- and *mTORC1*-targeted genes. High module scores suggest high expression of the gene set and are illustrated in red. Medium and low scores are illustrated in yellow and blue, respectively. The same color scheme is used for **A-D**. (**B**) UMAP projection of gene expression signatures of light zone (LZ), dark zone (DZ), and memory B cells defined among murine HA$^+$ B cells post LCMV infection by Mathew et al. (**C**) UMAP projection of gene expression signatures of Plasmablast, memory B cell (MBC), and GC B cells defined among human tonsillar B cells by King et al. (**D**) UMAP projection of gene expression signatures of LZ and DZ GC B cells defined among human tonsillar B cells by Holmes et al. (**E**) UMAP projection of cell latent time. Yellow color represents high latent time and dark blue represents low latent time. (**F**) The fraction of cells in each phenotypic cluster per mouse. The error bars represent the standard error of the mean. (**G**)

Violin plot of expression levels of *Myc, mTORC1,* and *E2f* separated by clusters. For (**F-G**), p values are computed with Kruskal-Wallis analysis of variance.

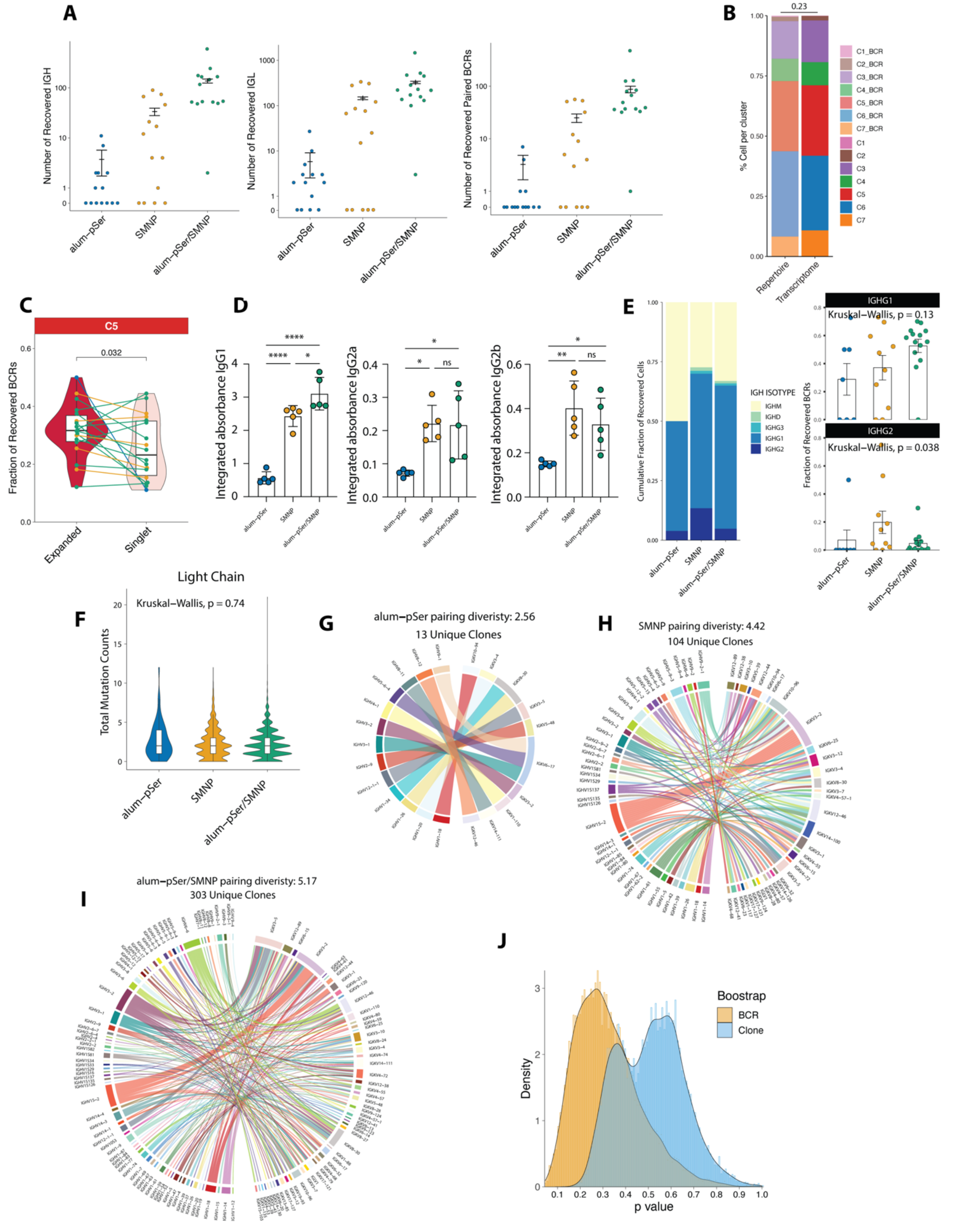

**Fig. S3. BCR repertoire of MD39-binding GC B cells.** (**A**) The number of heavy chain, light chain, and paired BCR sequences recovered from mice. Shown are mean ± s.e.m. (**B**) Cluster distribution of cells with paired BCR sequences recovered and all recovered cells. (**C**) Fraction of C5 cells per mouse from expanded clones vs. singlet clones. Only mice with both expanded and singlet clones recovered for the clusters were included in the analysis. The colored lines represent individual mice. The alum-pSer group is illustrated in blue, SMNP in orange, and alum-pSer/SMNP in green. P values were computed using permutation tests (100,000 trials). (**D**) BALB/c mice (*n*=5/group) were immunized with 5 µg MD39 ± 50 µg alum ± 5 µg SMNP. Serum IgG1, IgG2a, and IgG2b antibody responses were assessed at day 28 by ELISA using MD39 captured by lectin. Values plotted are the ELISA area under the curve (AUC) mean ± standard deviation. Statistical significance was determined by one-way ANOVA followed by Tukey's multiple comparisons test. ns $p>0.05$, * $p<0.05$, ** $p<0.01$, *** $p<0.001$, **** $p<0.0001$. (**E**) Distribution of heavy chain isotypes among MD39-binding GC B cells, and fractions of Ighg1 and Ighg2 BCRs per mouse. Error bars are plotted as the standard error of the mean. (**F**) Violin plots showing the light chain nucleotide mutation. (**G-I**) Chord diagram illustrating the clonal heavy and light chain pairings. Each chord on the diagram represents one clone, and the chord color represents the heavy chain V gene. Pairing diversity was calculated by the Shannon diversity index. (**J**) 200 BCRs or 100 clones were randomly sampled from the alum-pSer/SMNP ($N_{BCR}$=1222, $N_{clone}$=303) and SMNP ($N_{BCR}$=225, $N_{clone}$=104) groups, pairing diversity scores were calculated for each mouse, and p-value was computed with two-tailed t-test. This process was repeated 10,000 times to generate distributions of p-values.

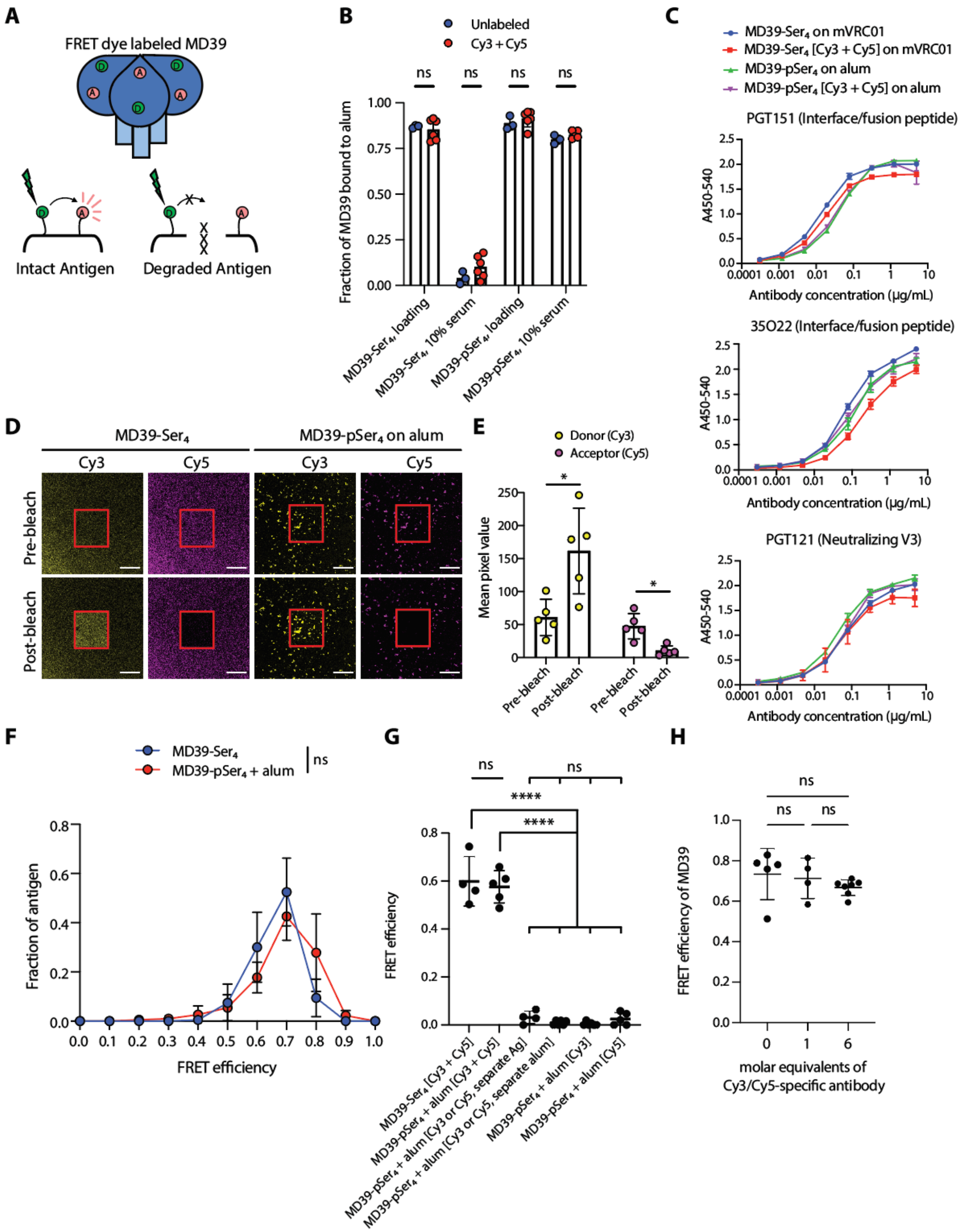

**Fig. S4. FRET-based approach allows for the assessment of antigen stability**. (**A**) Overview of FRET-based approach to investigate MD39 antigen stability. MD39 is labeled with donor and acceptor FRET pair dyes Cy3 and Cy5, such that FRET occurs when antigen is intact, and FRET does not occur when antigen is degraded. (**B**) pSer- and Ser-conjugated MD39 trimers were mixed with alum, and the fraction of protein bound to alum was assessed after loading and after 24-hour incubation in 10% mouse serum at 37°C. Statistical significance was determined by two-way ANOVA followed by Sidak's multiple comparisons test. (**C**) Antigenicity profiling of unlabeled and Cy3 + Cy5-labeled MD39 trimers. MD39-pSer$_4$ was captured by alum and MD39-Ser$_4$ was captured by mVRC01. Values plotted are means ± standard deviation. (**D**) Acceptor (Cy5) photobleaching approach for soluble and alum anchored MD39 coated on glass coverslips. Acceptor photobleaching of intact antigen labeled with Cy3 and Cy5 results in a reduction in Cy5 emission and an increase in Cy3 emission, quantified in (**E**) for soluble MD39. Scale bar represents 50 μm. Red outline indicates region of photobleaching. Statistical significance was determined by paired one-way ANOVA followed by Tukey's multiple comparisons test. (**F**) Histogram of FRET efficiencies (*n*=4-5). Values plotted are means ± standard deviation. Statistical significance was determined by unpaired Student's t-test. (**G**) FRET efficiencies of indicated proteins coated on glass coverslips. Values plotted are means ± standard deviation. Statistical significance was determined by one-way ANOVA followed by Tukey's multiple comparisons test. (**H**) MD39 was mixed with the indicated molar equivalents of a Cy3/Cy5-specific antibody and coated on glass coverslips. FRET efficiency values plotted are means ± standard deviation. Statistical significance was determined by one-way ANOVA followed by Tukey's multiple comparisons test. ns p>0.05. ns p>0.05, * p<0.05, ** p<0.01, *** p<0.001, **** p<0.0001.

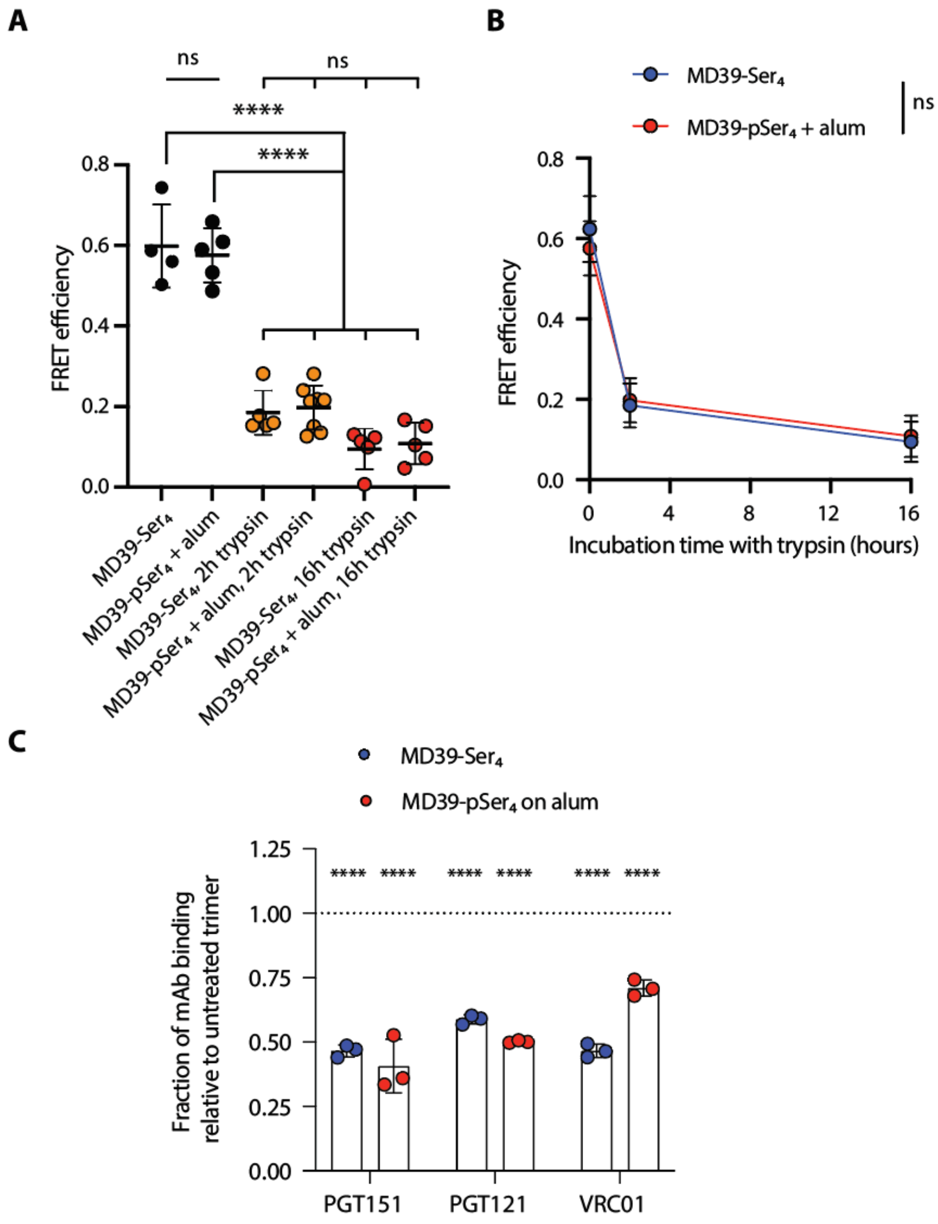

**Fig. S5. FRET efficiency is correlated to MD39 stability.** (**A**) FRET efficiencies of soluble MD39-Ser$_4$ and MD39-pSer$_4$ on alum, coated on glass coverslips following incubation with trypsin. Shown as longitudinal plot in (**B**). Values plotted are means ± standard deviation. Statistical significance was determined by two-way ANOVA followed by Tukey's multiple comparisons test. (**C**) Antigenicity profiling of soluble MD39-Ser$_4$ captured by 12N antibody and MD39-pSer$_4$ captured on alum following incubation with trypsin for 2 hours at 37°C. Shown are integrated absorbance values following incubation with trypsin for 2 hours at 37°C normalized to signal not exposed to trypsin. Values plotted are means ± standard deviation. Statistical significance was determined relative to the pre-2h incubation with trypsin values by two-way ANOVA followed by Tukey's multiple comparisons test. ns $p>0.05$, * $p<0.05$, ** $p<0.01$, *** $p<0.001$, **** $p<0.0001$.

**Data S1: Myc and mTORC1 target genes.**

| | Target Genes |
|---|---|
| **SCHUHMACHER_MYC_TARGETS_UP** | Abce1, Acsl1, Ahcy, Aimp2, Ak4, Akap1, Atp1b3, Auh, Bop1, Cad, Cdk4, Cebpz, Ctps1, Ctsc, Cycs, Cyp51, Dancr, Dcun1d4, Ddx10, Ddx21, Dhodh, Ebna1bp2, Exosc2, Exosc7, Fabp5, Fasn, Fkbp4, Fxn, Gcsh, Gpd1l, Grsf1, Hspe1, Iars1, Impdh2, Ldha, Lrp8, Mest, Mgst1, Mrpl3, Mthfd1, Mxi1, Myc, Nampt, Nefh, Nme1, Nolc1, Odc1, Paics, Pebp1, Pno1, Pold2, Polr2h, Ppat, Prdx4, Prps2, Pum3, Pycr1, Rabepk, Ranbp1, Rcc1, Rpia, Rrp1b, Rrs1, Slc16a1, Slc20a1, Slc39a14, Slc39a6, Sord, Srm, Srpk1, Tarbp1, Tbl3, Tfrc, Tmem97, Trap1, Uchl3, Uck2, Vars1, Vrk1, Znf239 |
| **PENG_RAPAMYCIN_RESPONSE_DN** | Adrm1, Ahsa1, Aldh4a1, Atf5, Atp5g1, Atp5g3, Bmp1, Btk, Bysl, C8orf41, Calu, Cct5, Cct6a, Cdk4, Chuk, Copb2, Cse1l, Eif2b4, Eif4g2, Emp3, Fam89b, Fasn, Hccs, Hmhb1, Hspa4, Hspa8, Hspe1, Hyou1, Id2, Ifi30, Imp4, Jtb, Lrrc32, Mllt11, Mmp15, Mrpl12, Mrps11, Ndufa9, Npm3, Pfkm, Pgk1, Prmt1, Psmd5, Psmd8, Ptbp1, Rabepk, Rabggtb, Safb, Sf3a3, Sit1, Slc29a1, Snrpc, Sqle, Srm, Tceb3, Tfrc, Tgds, Timm17a, Tmed2, Tmem109, Ube2l3, Wnt10b, Znf259 |

**Table S1: SeqWell primers.**

| **Primer Name** | **IDT Order** |
|---|---|
| SeqWell_WTA_Primer (TSO_PCR): | AAGCAGTGGTATCAACGCAGAGT |
| HTO_WTA_Primer | GTGACTGGAGTTCAGACGTGTGCTCTTCCGATCT |

**Table S2: Lockdown primers.**

| Primer Name | IDT Order |
|---|---|
| IGHM_mouse_lockdown | /5Biosg/CCTTCCCAAATGTCTTCCCCCTCGTCTCCTGCGAGAGCC CCCTGTCTGATAAGAATCTGGTGGCCATGGGCTGCCTIGCCCGG GACTTCC |
| IGHD_mouse_lockdown | /5Biosg/GAAATCCCACCATCTACCCACTGACACTCCCACGAGCT CTGTCAAGTGACCCAGTGATAATCGGCTGCCTGATTCACGATT ACTTCCCTT |
| IGHA_balb_lockdown | /5Biosg/GTGATAATCGGCTGCCTGATTCACGATTACTTCCCTTIC GGCACGATGAATGTGACCTGGGGAAAGAGTGGGAAGGATATA ACCACCGT |
| IGHG12AB_balb_lockdown | /5Biosg/TGCCTGGTCAAGGGITAITTCCCTGAGCCAGTGACIITGA CITGGAACTCTGGATCCCTGTCCAGIIGTGTGCACACCTTCCCAG CTITC |
| IGHG3_balb_lockdown | /5Biosg/CAGCCCCATCTGTCTATCCCTTGGTCCCTGGCTGCGGTG ACACATCTGGATCCTCGGTGACACTGGGATGCCTTGTCAAAGG CTACTTCC |
| IGKC_mouse_lockdown | /5Biosg/TGCACCAACTGTATCCATCTTCCCACCATCCAGTGAGCA GTTAACATCTGGAGGTGCCTCAGTCGTGTGCTTCTTGAACAACT TCTACCC |
| IGLC_mouse_lockdown | /5Biosg/GTCTTCGCCATCAGTCACCCTGTTTCCACCTTCCTCTGAA GAGCTCGAGACTAACAAGGCCACACTGGTGTGTACGATCACTGA TTTCTA |
| IGLC23_mouse_lockdown | /5Biosg/AGTGGTGTGACAGTGGCCTGGAAGGCAAATGGTACACC TATCACCCAGGGTGTGGACACTTCAAATCCCACCAAAGAGGGC AACAAGTTC |

**Table S3: V gene primers.**

| Primer Name | IDT Order | Nextera Seq Handle | 5'-3' Seq |
|---|---|---|---|
| mVH01 | TCGTGGGCTCGGAGATGTGTATAAGAGACAGCAGGTGCAGCTGCAGCAGCCTGG | TCGTGGGCTCGGAGATGTGTATAAGAGACAG | CAGGTGCAGCTGCAGCAGCCTGG |
| mVH02 | TCGTGGGCTCGGAGATGTGTATAAGAGACAGCAGGTGCAGCTGCAGCAGTCTGG | TCGTGGGCTCGGAGATGTGTATAAGAGACAG | CAGGTGCAGCTGCAGCAGTCTGG |
| mVH03 | TCGTGGGCTCGGAGATGTGTATAAGAGACAGCAGGTGCAGCTGAAGCAGTCTGG | TCGTGGGCTCGGAGATGTGTATAAGAGACAG | CAGGTGCAGCTGAAGCAGTCTGG |
| mVH04 | TCGTGGGCTCGGAGATGTGTATAAGAGACAGCAGGTGCAGCTGAAGGAGTCTGG | TCGTGGGCTCGGAGATGTGTATAAGAGACAG | CAGGTGCAGCTGAAGGAGTCTGG |
| mVH05 | TCGTGGGCTCGGAGATGTGTATAAGAGACAGGAGGTGAAGCTGGAGGAGTCTGG | TCGTGGGCTCGGAGATGTGTATAAGAGACAG | GAGGTGAAGCTGGAGGAGTCTGG |
| mVH06 | TCGTGGGCTCGGAGATGTGTATAAGAGACAGGAGGTGCAGCTGGTGGAGTCTGG | TCGTGGGCTCGGAGATGTGTATAAGAGACAG | GAGGTGCAGCTGGTGGAGTCTGG |
| mVH07 | TCGTGGGCTCGGAGATGTGTATAAGAGACAGGAAGTGCAGCTGTTGGAGACTGG | TCGTGGGCTCGGAGATGTGTATAAGAGACAG | GAAGTGCAGCTGTTGGAGACTGG |
| mVH08 | TCGTGGGCTCGGAGATGTGTATAAGAGACAGGAGGTGCAGCTGCAGCAGTCTGG | TCGTGGGCTCGGAGATGTGTATAAGAGACAG | GAGGTGCAGCTGCAGCAGTCTGG |
| mVH09 | TCGTGGGCTCGGAGATGTGTATAAGAGACAGGAGGTGCAGCTGCAGGAGTCTGG | TCGTGGGCTCGGAGATGTGTATAAGAGACAG | GAGGTGCAGCTGCAGGAGTCTGG |
| mVH10 | TCGTGGGCTCGGAGATGTGTATAAGAGACAGGAGGTGCAGCTGCAGCAGTCTGTG | TCGTGGGCTCGGAGATGTGTATAAGAGACAG | GAGGTGCAGCTGCAGCAGTCTGTG |
| mVH11 | TCGTGGGCTCGGAGATGTGTATAAGAGACAGGAGGTGAAGCTGGTGGAGTCTGG | TCGTGGGCTCGGAGATGTGTATAAGAGACAG | GAGGTGAAGCTGGTGGAGTCTGG |
| mVH12 | TCGTGGGCTCGGAGATGTGTATAAGAGACAGCAGATCCAGCTGCAGCAGTCTGG | TCGTGGGCTCGGAGATGTGTATAAGAGACAG | CAGATCCAGCTGCAGCAGTCTGG |
| mVH13 | TCGTGGGCTCGGAGATGTGTATAAGAGACAGCAGGTTCAGCTGCAACAGTCTGA | TCGTGGGCTCGGAGATGTGTATAAGAGACAG | CAGGTTCAGCTGCAACAGTCTGA |
| mVH14 | TCGTGGGCTCGGAGATGTGTATAAGAGACAGGAGTTCCAGCTGCAGCAGTCTGG | TCGTGGGCTCGGAGATGTGTATAAGAGACAG | GAGTTCCAGCTGCAGCAGTCTGG |
| mVH15 | TCGTGGGCTCGGAGATGTGTATAAGAGACAGGATGTACAGCTTCAGGAGTCAGG | TCGTGGGCTCGGAGATGTGTATAAGAGACAG | GATGTACAGCTTCAGGAGTCAGG |
| mVH16 | TCGTGGGCTCGGAGATGTGTATAAGAGACAGGAGGTGCAGCTTGTTGAGTCTGGTGGA | TCGTGGGCTCGGAGATGTGTATAAGAGACAG | GAGGTGCAGCTTGTTGAGTCTGGTGGAGG |

| | | | |
|---|---|---|---|
| | GG | | |
| mVH17 | TCGTGGGCTCGGAGATGTG TATAAGAGACAGCAGCGTG AGCTGCAGCAGTCTGG | TCGTGGGCTCGGAGA TGTGTATAAGAGACAG | CAGCGTGAGC TGCAGCAGTC TGG |
| mVH18 | TCGTGGGCTCGGAGATGTG TATAAGAGACAGGACGTGA AGCTGGTGGAGTCTGG | TCGTGGGCTCGGAGA TGTGTATAAGAGACAG | GACGTGAAGC TGGTGGAGTC TGG |
| mVH19 | TCGTGGGCTCGGAGATGTG TATAAGAGACAGGAAGTGA TGCTGGTGGAGTCTGG | TCGTGGGCTCGGAGA TGTGTATAAGAGACAG | GAAGTGATGC TGGTGGAGTC TGG |
| mVH20 | TCGTGGGCTCGGAGATGTG TATAAGAGACAGCAGGTGC AGCTTGTAGAGACCGG | TCGTGGGCTCGGAGA TGTGTATAAGAGACAG | CAGGTGCAGC TTGTAGAGAC CGG |
| mVH21 | TCGTGGGCTCGGAGATGTG TATAAGAGACAGCAGATGC AGCTTCAGGAGTCAGG | TCGTGGGCTCGGAGA TGTGTATAAGAGACAG | CAGATGCAGC TTCAGGAGTC AGG |
| mVH22 | TCGTGGGCTCGGAGATGTG TATAAGAGACAGCAGGCTT ATCTACAGCAGTCTGG | TCGTGGGCTCGGAGA TGTGTATAAGAGACAG | CAGGCTTATC TACAGCAGTC TGG |
| mVH23 | TCGTGGGCTCGGAGATGTG TATAAGAGACAGCAGGTCC ARCTGCAGCAGYCTGG | TCGTGGGCTCGGAGA TGTGTATAAGAGACAG | CAGGTCCARC TGCAGCAGYC TGG |
| mVH24 | TCGTGGGCTCGGAGATGTG TATAAGAGACAGGAGGTGA AGCTTCTCSAGTCTGGAGG | TCGTGGGCTCGGAGA TGTGTATAAGAGACAG | GAGGTGAAGC TTCTCSAGTCT GGAGG |
| mVH25 | TCGTGGGCTCGGAGATGTG TATAAGAGACAGCAGGTTA CTCTGAAAGAGTCTGGCC | TCGTGGGCTCGGAGA TGTGTATAAGAGACAG | CAGGTTACTC TGAAAGAGTC TGGCC |
| mVH26 | TCGTGGGCTCGGAGATGTG TATAAGAGACAGCAGGGTC AGATGCAGCAGTCTGG | TCGTGGGCTCGGAGA TGTGTATAAGAGACAG | CAGGGTCAGA TGCAGCAGTC TGG |
| mVK01 | TCGTGGGCTCGGAGATGTG TATAAGAGACAGAACATTA TGATGACACAGTCGCCA | TCGTGGGCTCGGAGA TGTGTATAAGAGACAG | AACATTATGA TGACACAGTC GCCA |
| mVK02 | TCGTGGGCTCGGAGATGTG TATAAGAGACAGAACATTG TGCTGACCCAATCTCCA | TCGTGGGCTCGGAGA TGTGTATAAGAGACAG | AACATTGTGC TGACCCAATC TCCA |
| mVK03 | TCGTGGGCTCGGAGATGTG TATAAGAGACAGCAAATTG TTCTCACCCAGTCTCCA | TCGTGGGCTCGGAGA TGTGTATAAGAGACAG | CAAATTGTTC TCACCCAGTC TCCA |
| mVK04 | TCGTGGGCTCGGAGATGTG TATAAGAGACAGCAAATTG TTCTCTCCCAGTCTCCA | TCGTGGGCTCGGAGA TGTGTATAAGAGACAG | CAAATTGTTC TCTCCCAGTC TCCA |
| mVK05 | TCGTGGGCTCGGAGATGTG TATAAGAGACAGGAAAATG TTCTCACCCAGTCTCCA | TCGTGGGCTCGGAGA TGTGTATAAGAGACAG | GAAAATGTTC TCACCCAGTC TCCA |
| mVK06 | TCGTGGGCTCGGAGATGTG TATAAGAGACAGGAAATTG | TCGTGGGCTCGGAGA TGTGTATAAGAGACAG | GAAATTGTGC TCACTCAGTC |

| | | | |
|---|---|---|---|
| | TGCTCACTCAGTCTCCA | | TCCA |
| mVK07 | TCGTGGGCTCGGAGATGTG TATAAGAGACAGGACATCA AGATGACCCAGTCTCCA | TCGTGGGCTCGGAGA TGTGTATAAGAGACAG | GACATCAAGA TGACCCAGTC TCCA |
| mVK08 | TCGTGGGCTCGGAGATGTG TATAAGAGACAGGACATCC AGATGAACCAGTCTCCA | TCGTGGGCTCGGAGA TGTGTATAAGAGACAG | GACATCCAGA TGAACCAGTC TCCA |
| mVK09 | TCGTGGGCTCGGAGATGTG TATAAGAGACAGGACATCC AGATGACTCAGTCTCCA | TCGTGGGCTCGGAGA TGTGTATAAGAGACAG | GACATCCAGA TGACTCAGTC TCCA |
| mVK10 | TCGTGGGCTCGGAGATGTG TATAAGAGACAGGACATTG TGATGACTCAGTCTC | TCGTGGGCTCGGAGA TGTGTATAAGAGACAG | GACATTGTGA TGACTCAGTC TC |
| mVK11 | TCGTGGGCTCGGAGATGTG TATAAGAGACAGGACATTG TGATGTCACAGTCTCCA | TCGTGGGCTCGGAGA TGTGTATAAGAGACAG | GACATTGTGA TGTCACAGTC TCCA |
| mVK12 | TCGTGGGCTCGGAGATGTG TATAAGAGACAGGACATTG TGCTGACCCAATCTCCA | TCGTGGGCTCGGAGA TGTGTATAAGAGACAG | GACATTGTGC TGACCCAATC TCCA |
| mVK13 | TCGTGGGCTCGGAGATGTG TATAAGAGACAGGATATCC AGATGACACAGACTACA | TCGTGGGCTCGGAGA TGTGTATAAGAGACAG | GATATCCAGA TGACACAGAC TACA |
| mVK14 | TCGTGGGCTCGGAGATGTG TATAAGAGACAGGATGTTG TGATGACCCAAACTCCA | TCGTGGGCTCGGAGA TGTGTATAAGAGACAG | GATGTTGTGA TGACCCAAAC TCCA |
| mVK15 | TCGTGGGCTCGGAGATGTG TATAAGAGACAGGAAATCC AGATGACCCAGTCTCCA | TCGTGGGCTCGGAGA TGTGTATAAGAGACAG | GAAATCCAGA TGACCCAGTC TCCA |
| mVK16 | TCGTGGGCTCGGAGATGTG TATAAGAGACAGGACATCC AGATGACACAATCTTCA | TCGTGGGCTCGGAGA TGTGTATAAGAGACAG | GACATCCAGA TGACACAATC TTCA |
| mVK17 | TCGTGGGCTCGGAGATGTG TATAAGAGACAGGACATCC AGATGACCCAGTCTCCA | TCGTGGGCTCGGAGA TGTGTATAAGAGACAG | GACATCCAGA TGACCCAGTC TCCA |
| mVK18 | TCGTGGGCTCGGAGATGTG TATAAGAGACAGGACATCC TGATGACCCAATCTCCA | TCGTGGGCTCGGAGA TGTGTATAAGAGACAG | GACATCCTGA TGACCCAATC TCCA |
| mVK19 | TCGTGGGCTCGGAGATGTG TATAAGAGACAGGACATTG TGCTCACCCAATCTCC | TCGTGGGCTCGGAGA TGTGTATAAGAGACAG | GACATTGTGC TCACCCAATC TCC |
| mVK20 | TCGTGGGCTCGGAGATGTG TATAAGAGACAGGATGTTG TGGTGACTCAAACTCCA | TCGTGGGCTCGGAGA TGTGTATAAGAGACAG | GATGTTGTGG TGACTCAAAC TCCA |
| mVK21 | TCGTGGGCTCGGAGATGTG TATAAGAGACAGAACATTG TAATGACCCAATCTCCC | TCGTGGGCTCGGAGA TGTGTATAAGAGACAG | AACATTGTAA TGACCCAATC TCCC |
| mVK22 | TCGTGGGCTCGGAGATGTG TATAAGAGACAGGATGTTT TGATGACCCAAACTCCA | TCGTGGGCTCGGAGA TGTGTATAAGAGACAG | GATGTTTTGA TGACCCAAAC TCCA |

| | | | |
|---|---|---|---|
| mVK23 | TCGTGGGCTCGGAGATGTG TATAAGAGACAGGACATCC AGATGATTCAGTCTCCA | TCGTGGGCTCGGAGA TGTGTATAAGAGACAG | GACATCCAGA TGATTCAGTC TCCA |
| mVK24 | TCGTGGGCTCGGAGATGTG TATAAGAGACAGGACATCT TGCTGACTCAGTCTCCA | TCGTGGGCTCGGAGA TGTGTATAAGAGACAG | GACATCTTGC TGACTCAGTC TCCA |
| mVK25 | TCGTGGGCTCGGAGATGTG TATAAGAGACAGGATGTCC AGATGATTCAGTCTCCA | TCGTGGGCTCGGAGA TGTGTATAAGAGACAG | GATGTCCAGA TGATTCAGTC TCCA |
| mVK26 | TCGTGGGCTCGGAGATGTG TATAAGAGACAGGATGTCC AGATAACCCAGTCTCCA | TCGTGGGCTCGGAGA TGTGTATAAGAGACAG | GATGTCCAGA TAACCCAGTC TCCA |
| mVK27 | TCGTGGGCTCGGAGATGTG TATAAGAGACAGGACATTG TGATGACCCAGTCTCAM | TCGTGGGCTCGGAGA TGTGTATAAGAGACAG | GACATTGTGA TGACCCAGTC TCAM |
| mVL01 | TCGTGGGCTCGGAGATGTG TATAAGAGACAGRGCTGTT GTGACTCAGGAATC | TCGTGGGCTCGGAGA TGTGTATAAGAGACAG | RGCTGTTGTG ACTCAGGAA TC |

**Table S4: Index PCR primers.**

| Primer Name | IDT Order | Index Name | P7 | Index | Adapter |
|---|---|---|---|---|---|
| P7_index_Next | CAAGCAGAAGACGGCATACGAGATGCAGCGTATCGTGGGCTCGGAGATGTG | N721 | CAAGCAGAAGACGGCATACGAGAT | GCAGCGTA | TCGTGGGCTCGGAGATGTG |
| P5_index_TSO01 | AATGATACGGCGACCACCGAGATCTACACTAGATCGCGCCTGTCCGCGGAAGCAGTG | N501 | AATGATACGGCGACCACCGAGATCTACAC | TAGATCGC | GCCTGTCCGCGGAAGCAGTG |
| P5_index_TSO02 | AATGATACGGCGACCACCGAGATCTACACCTCTCTATGCCTGTCCGCGGAAGCAGTG | N502 | AATGATACGGCGACCACCGAGATCTACAC | CTCTCTAT | GCCTGTCCGCGGAAGCAGTG |
| P5_index_TSO03 | AATGATACGGCGACCACCGAGATCTACACTATCCTCTGCCTGTCCGCGGAAGCAGTG | N503 | AATGATACGGCGACCACCGAGATCTACAC | TATCCTCT | GCCTGTCCGCGGAAGCAGTG |
| P5_index_TSO04 | AATGATACGGCGACCACCGAGATCTACACAGAGTAGAGCCTGTCCGCGGAAGCAGTG | N504 | AATGATACGGCGACCACCGAGATCTACAC | AGAGTAGA | GCCTGTCCGCGGAAGCAGTG |
| P5_index_TSO05 | AATGATACGGCGACCACCGAGATCTACACGTAAGGAGGCCTGTCCGCGGAAGCAGTG | N505 | AATGATACGGCGACCACCGAGATCTACAC | GTAAGGAG | GCCTGTCCGCGGAAGCAGTG |
| P5_index_TSO06 | AATGATACGGCGACCACCGAGATCTACACACTGCATAGCCTGTCCGCGGAAGCAGTG | N506 | AATGATACGGCGACCACCGAGATCTACAC | ACTGCATA | GCCTGTCCGCGGAAGCAGTG |
| P5_index_TSO07 | AATGATACGGCGACCACCGAGATCTACACCTAAGCCTGCCTGTCCGCGGAAGCAGTG | N508 | AATGATACGGCGACCACCGAGATCTACAC | CTAAGCCT | GCCTGTCCGCGGAAGCAGTG |
| P5_index_TSO08 | AATGATACGGCGACCACCGAGATCTACACCGTCTAATGCCTGTCCGCGGAAGCAGTG | N510 | AATGATACGGCGACCACCGAGATCTACAC | CGTCTAAT | GCCTGTCCGCGGAAGCAGTG |
| P5_index_TSO09 | AATGATACGGCGACCACCGAGATCTACACTCTCTCCGGCCTGTCCGCGGAAGCAGTG | N511 | AATGATACGGCGACCACCGAGATCTACAC | TCTCTCCG | GCCTGTCCGCGGAAGCAGTG |
| P5_index_TSO10 | AATGATACGGCGACCACCGAGATCTACACTCGACTAGGCCTGTCCGCGGAAGCAGTG | N513 | AATGATACGGCGACCACCGAGATCTACAC | TCGACTAG | GCCTGTCCGCGGAAGCAGTG |
| P5_index_TSO11 | AATGATACGGCGACCACCGAGATCTACACT | N515 | AATGATACGGCGACCACCGAGA | TTCTAGCT | GCCTGTCCGCGGAA |

| | | | | | |
|---|---|---|---|---|---|
| | TCTAGCTGCCTGTCC GCGGAAGCAGTG | | TCTACAC | | GCAGTG |
| P5_index_TSO12 | AATGATACGGCGACC ACCGAGATCTACACC CTAGAGTGCCTGTCC GCGGAAGCAGTG | N516 | AATGATACGGC GACCACCGAGA TCTACAC | CCTAGAGT | GCCTGTC CGCGGAA GCAGTG |
| P5_index_TSO13 | AATGATACGGCGACC ACCGAGATCTACACG CGTAAGAGCCTGTCC GCGGAAGCAGTG | N517 | AATGATACGGC GACCACCGAGA TCTACAC | GCGTAAGA | GCCTGTC CGCGGAA GCAGTG |
| P5_index_TSO14 | AATGATACGGCGACC ACCGAGATCTACACA AGGCTATGCCTGTCC GCGGAAGCAGTG | N520 | AATGATACGGC GACCACCGAGA TCTACAC | AAGGCTAT | GCCTGTC CGCGGAA GCAGTG |
| P5_index_TSO15 | AATGATACGGCGACC ACCGAGATCTACACG AGCCTTAGCCTGTCC GCGGAAGCAGTG | N521 | AATGATACGGC GACCACCGAGA TCTACAC | GAGCCTTA | GCCTGTC CGCGGAA GCAGTG |
| P5_index_TSO16 | AATGATACGGCGACC ACCGAGATCTACACT TATGCGAGCCTGTCC GCGGAAGCAGTG | N522 | AATGATACGGC GACCACCGAGA TCTACAC | TTATGCGA | GCCTGTC CGCGGAA GCAGTG |

**Table S5: Sequencing primers.**

| Primer Name | IDT Order |
|---|---|
| IGHM_mouse_seq | GCTCTCGCAGGAGACGAGGGGGAAGACATTTGGGAA |
| IGHD_mouse_seq | GGGCTTTGCACTCTGAGAGGAGGAACATGTCAG |
| IGHG12AB_mouse_seq | CAGGGGCCAGTGGATAGACIGATGGGGITGT |
| IGHG3_mouse_seq | CCGAGGATCCAGATGTGTCACCGCAGCCAGGG |
| IGHA_mouse_seq | AGGACTGGTGGGAGTGTCAGTGGGTAGATGGTGGGAT |
| IGLC1_mouse_seq | TCTTCAGAGGAAGGTGGAAACAGGGTGACTGATGGCGAA |
| IGLC2_mouse_seq | CTCAGAGGAAGGTGGAAACAIGGTGAGIGTGGGAGTGG |
| IGKC_mouse_seq | TCCAGATGTTAACTGCTCACTGGATGGTGGGAAGATGGA TACAGTTGGTG |
| SeqWell Read 1 | GCCTGTCCGCGGAAGCAGTGGTATCAACGCAGAGTAC |
| SeqWell_Index_Read2_primer | ACTCTGCGTTGATACCACTGCTTCCGCGGACAGGC |
